# Generation and characterisation of an *Arabidopsis thaliana f3h*/*fls1*/*ans* triple mutant that accumulates eriodictyol derivatives

**DOI:** 10.1101/2023.09.21.558826

**Authors:** Hanna Marie Schilbert, Mareike Busche, Vania Sáez, Andrea Angeli, Bernd Weisshaar, Stefan Martens, Ralf Stracke

## Abstract

**Background:** Flavonoids are plant specialised metabolites, which derive from phenylalanine and acetate metabolism. They possess a variety of beneficial characteristics for plants and humans. Several modification steps in the synthesis of tricyclic flavonoids cause for the amazing diversity of flavonoids in plants. The 2-oxoglutarate-dependent dioxygenases (2-ODDs) flavanone 3-hydroxylase (F3H, synonym FHT), flavonol synthase (FLS) and anthocyanidin synthase (ANS, synonym leucoanthocyanidin dioxigenase (LDOX)), catalyse oxidative modifications to the central C ring. They are highly similar and have been shown to catalyse, at least in part, each other’s reactions. FLS and ANS have been identified as bifunctional enzymes in many species, including *Arabidopsis thaliana*, stressing the capability of plants to bypass missing or mutated reaction steps on the way to flavonoid production. However, little is known about such bypass reactions and the flavonoid composition of plants lacking all three central flavonoid 2-ODDs.

**Results:** To address this issue, we generated a *f3h*/*fls1*/*ans* mutant, as well as the corresponding double mutants and investigated the flavonoid composition of this mutant collection. The *f3h*/*fls1*/*ans* mutant was further characterised at the genomic level by analysis of a nanopore DNA sequencing generated genome sequence assembly and at the transcriptomic level by RNA-Seq analysis. The mutant collection established, including the novel double mutants *f3h*/*fls1* and *f3h*/*ans*, was used to validate and analyse the multifunctionalities of F3H, FLS1, and ANS *in planta*. Metabolite analyses revealed the accumulation of eriodictyol and additional glycosylated derivatives in mutants carrying the *f3h* mutant allele, resulting from the conversion of naringenin to eriodictyol by flavonoid 3’-hydroxylase (F3’H) activity.

****Conclusions**:** We describe the *in planta* multifunctionality of the three central flavonoid 2-ODDs from *A. thaliana* and identify a bypass in the *f3h*/*fls1*/*ans* triple mutant that leads to the formation of eriodictyol derivatives. As (homo-)eriodictyols are known as bitter taste maskers, the annotated eriodictyol (derivatives) and in particular the observations made on their *in planta* production, could provide valuable insights for creating of novel food supplements.

## Background

Flavonoids are a large group of specialised metabolites with an estimated number of 9,000 flavonoid derivatives and a wide distribution throughout the plant kingdom (1). Flavonoids share a basic structure with two aromatic C6 rings and a heterocyclic ring (2). Reorganisation or modification, such as oxidation or glycosylation, is mainly responsible for the wide variety of flavonoids. They can be classified into different subgroups including flavones, flavanones, flavonols, dihydroflavonols, flavandiols, anthocyanins, and proanthocyanidins (2). These subgroups exhibit manifold functions. Among others, anthocyanins promote pollination and seed dispersal by colouring flowers and fruits (3–5), flavonols play a role in auxin transport, plant fertility, and UV protection (6–8), and proanthocyanins serve as deterrent metabolites for defence against bird predators (9). Moreover, flavonoids are beneficial for human nutrition due to their antioxidative, antigenotoxic and anticarcinogenic properties (10, 11). These health-promoting effects have led to a strong interest in increasing the flavonoid content in plants (12, 13). However, as some flavonoids exhibit antinutritional properties, like a bitter-off taste (14), the engineering of flavonoid composition is of high relevance for the breeding of plants with desirable flavonoid profiles.

Flavonoid biosynthesis (Figure 1) is one of the best characterised pathways in plant specialised metabolism. Due to the visible phenotypes of respective mutants which often display a transparent testa resulting in yellow to pale-brown seed (15) and the large collections of T-DNA- and transposon-tagged lines for *Arabidopsis thaliana* (16–18), the genes involved in flavonoid biosynthesis are well studied. The flavonoid pathway is initiated by its key enzyme chalcone synthase (CHS), which condensates *p*-coumaroyl-CoA and three molecules of malonyl-CoA followed by cyclisation, leading to the formation of naringenin chalcone (19). A heterocyclic ring is introduced spontaneously or through the catalysis of chalcone isomerase (CHI), forming naringenin (20). Flavanone 3-hydroxylase (F3H or FHT) converts naringenin, a flavanone, to dihydrokaempferol, a dihydroflavonol, by hydroxylation at the C-3 position (21). In alternative branches, the flavonoid 3’-hydroxylase (F3’H) converts naringenin to eriodictyol and dihydrokaempferol to dihydroquercetin (22). Naringenin can also be converted to flavones by flavone synthaseI or II (FNSI or II) (23, 24). However, flavones were thought to be absent in *A. thaliana* until two derivatives (7-2″,3″-diacetylglucoside and pentamethoxydihydroxy flavone) were identified in Columbia-0 (Col-0) leaves (25, 26). DOWNY MILDEW RESISTANT6 (*Ath*DMR6), a 2-oxoglutarate-dependent dioxygenase (2-ODD), has been proposed as the enzyme responsible for their biosynthesis (26). The enzyme flavonol synthase (FLS) uses dihydroflavonols to form flavonols by introducing a double bond between C-2 and C-3 (27). Alternatively, dihydroflavonol 4-reductase (DFR) can reduce dihydroflavonols to the corresponding leucoanthocyanidins, thus competing with FLS for substrates (28, 29). Anthocyanidin synthase (ANS, also known as leucoanthocyanidin dioxygenase (LDOX)) converts leucoanthocyanidins to anthocyanidins (30). Anthocyanidins can be modified by glycosyltransferases (GTs) to form anthocyanins, or they can be converted to proanthocyanidins by anthocyanidin reductase (ANR) (31, 32). Besides anthocyanidins, other flavonoids like flavonols or flavanones can be modified by GTs to form the respective glycosides (33–35).

**Figure 1:**
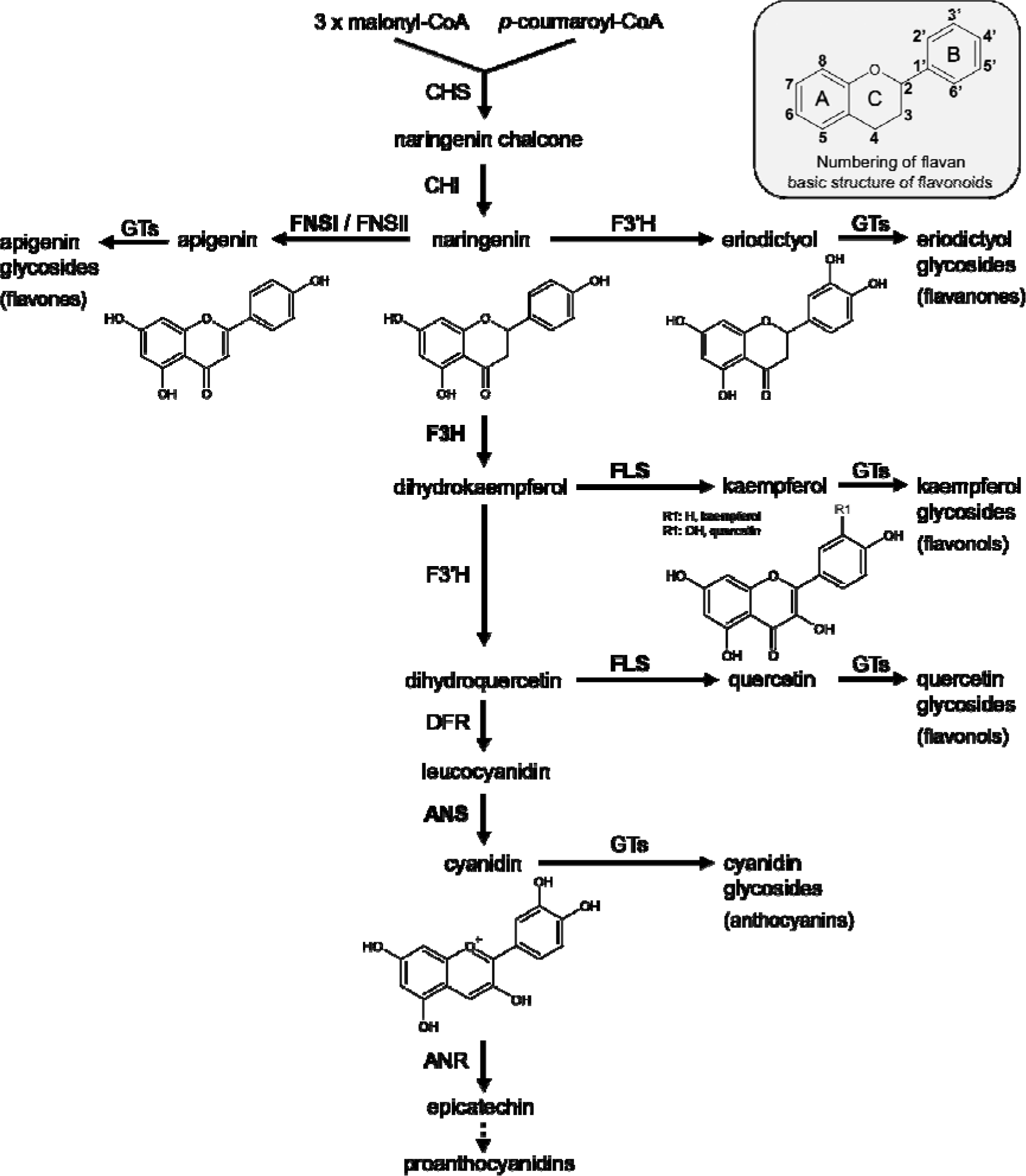
Simplified representation of the flavonoid biosynthesis pathway in plants. Details and abbreviations are given in the text. The 2-ODDs of the flavonoid biosynthesis F3H, FNSI, FLS, and ANS are shown in bold.

F3H, FNSI, FLS and ANS belong to the 2-ODD enzyme family (36). 2-ODDs represent the second largest enzyme family in plants (37). They are ubiquitously spread across the plant kingdom but also microorganisms and are involved in a variety of primary and specialised metabolism, including the synthesis of specialised metabolites, hormones, signalling molecules, amino acid modification and DNA-repair (summarised in 38, 39). Plant 2-ODDs can be classified into three classes, DOXA, DOXB and DOXC, based on amino acid sequence similarity, with DOXC harbouring 2-ODDs involved in the synthesis of specialised metabolites, such as flavonoids, including F3H and FNSI (DOXC28) and FLS and ANS (DOXC47) (37).

Characteristic hallmarks of 2-ODDs are generally (but not exclusively) the requirement of 2-oxoglutarate as co-reductant and iron as co-factor (36). Elucidation of the crystallographic structure of *Ath*ANS has identified the conserved amino acid residues involved in substrate-, co-factor-, and co-substrate binding (30). In addition, residues important for proper folding and substrate binding have been described (30).

As F3H, FLS and ANS all carry the conserved amino acid residues essential for 2-ODD activity, it has been proposed that they share a common catalytic mechanism, possibly allowing multifunctionality (40, 41). While FNSI and F3H often shows a narrow substrate spectrum (40, 42), FLS and ANS have been characterised as bi- or even multifunctional enzymes in several species (40, 43–46). From an evolutionary perspective, it seems feasible that the 2-ODD enzyme proposed to have evolved first, F3H, may have given rise to the other 2-ODDs involved in flavonoid biosynthesis (summarised in 47) and that *Ath*FLS1 and *Ath*ANS retain *in vitro* F3H (40, 48) and, in the case of *Ath*ANS, additional FLS side activity *in planta* (49). Due to the multifunctional nature of flavonoid 2-ODD enzymes, comprehensive biochemical and functional characterisation is required for unequivocal annotation, especially *in planta*. Only seven amino acid residues distinguish parsley F3H from FNSI, providing an impressive example that amino acid alignments are not sufficient for full functional characterisation (24).

The multifunctionalities of flavonoid 2-ODDs can be seen in the “leaky” phenotypes of *f3h*, *fls1* and *ans* mutants of *A. thaliana* (21, 27, 49, 50), attributed to the partial compensation by other 2-ODDs. The *ans* mutant shows a significant reduction in anthocyanin and proanthocyanidin content with almost unchanged flavonol levels compared to the wildtype (49, 51, 52), while the *fls1* mutant shows a significant reduction in flavonol content and increased anthocyanin and proanthocyanidin levels (49, 50, 53). The *f3h* mutant has a reduced flavonoid content, including flavonols, anthocyanins, and proanthocyanidins, due to a generally reduced flux in the flavonoid pathway (21, 54).

To unambiguously annotate the *in planta* multifunctionalities of F3H, FLS and ANS (which does not necessarily reflect the *in vitro* function), multiple mutants are required, like it was done in the case of the *fls1*/*ans* double mutant, which demonstrated the *in planta* FLS side activity of ANS (40, 49).

In this study, we generated novel *A. thaliana f3h*/*fls1* and *f3h*/*ans* double mutants, as well as an *f3h*/*fls1*/*ans* triple mutant to examine the phenotypic changes resulting from the loss of activity of these flavonoid 2-ODDs. The *f3h*/*fls1*/*ans* mutant was characterised in detail, using genomic, transcriptomic and metabolomic methods. Comprehensive phenotyping of the flavonoid composition of the established flavonoid 2-ODD mutant set (composed of single, double, and triple mutants) provided new insights into the *in planta* multifunctionalities of *A. thaliana* F3H, FLS1 and ANS and the resulting metabolic fluxes. We found a flux into a family of eriodictyol derivatives, which is most prominent in the *f3h*/*fls1*/*ans* triple mutant. The knowledge gained in this study will be valuable for targeted engineering of flavonoid composition and content in plants, especially crops.

## Results

### Genomic characterisation of the *A. thaliana f3h*/*fls1*/*ans* mutant

To generate an *A. thaliana f3h*/*fls1*/*ans* mutant set, we used previously published insertional alleles of the respective genes, showing the known loss-of-function phenotypes (*f3h* (*tt6-2*, GK-292E08, Col-0 background), *fls1* (*fls1-2,* RIKEN_PST16145, Nö-0 background), *ans* (*tds4-4*, SALK_073183, Col-0 background)). An *fls1*/*ans* double mutant was crossed with the *f3h* single mutant. By screening the F2 progeny, homozygous *f3h*/*fls1*/*ans*, *f3h*/*fls1* and *f3h*/*ans* mutants were identified by PCR-based genotyping. A *de novo* long-read assembly of the *f3h*/*fls1*/*ans* mutant genome was generated to analyse the respective mutated loci at the genomic level. The 160.57 Mbp assembly comprises 162 contigs with an N50 of 8.17 Mbp (Supplementary Table S1). The assembly size slightly exceeds the size of the Col-0 Community-Consensus reference genome sequence assembly (133,336,906 bp) likely due to the increased heterozygosity derived from the Nö-0 background of the *fls1* mutant. In total, 95.5 % complete BUSCOs were identified, indicating a high level of assembly completeness.

The assembly was used to validate the position of the T-DNA insertions at the *f3h* and *ans* loci, and the position of the immobilised Ds transposon at the *fls1* locus (Figure 2). Two T-DNAs comprising 11,393 bp sequence are inserted head-to-head into the 5’UTR of *f3h*, precisely between 19,025,263 - 19,025,278 bp on chromosome three. The 5,417 bp Ds transposon is inserted between 2,804,241 - 2,804,242 bp on chromosome five into the *fls1* locus in the first exon. A large T-DNA array comprised of several T-DNAs and the corresponding binary vector backbone (BVB) sequences was detected between 12,005,371 - 12,005,393 bp on chromosome four, inserted into the front part of the second exon of *ans*. Due to the high amount of repetitive sequences derived from this complex T-DNA array, the array is split across two contigs in the assembly and therefore the exact size (spans >119 kbp) could not be determined. A T-DNA left border (LB) was located at both ends of this insertion. Two additional T-DNA insertion events have been identified (Supplementary Figure S1). However, these insertions do not disrupt any additional annotated genes.

**Figure 2:**
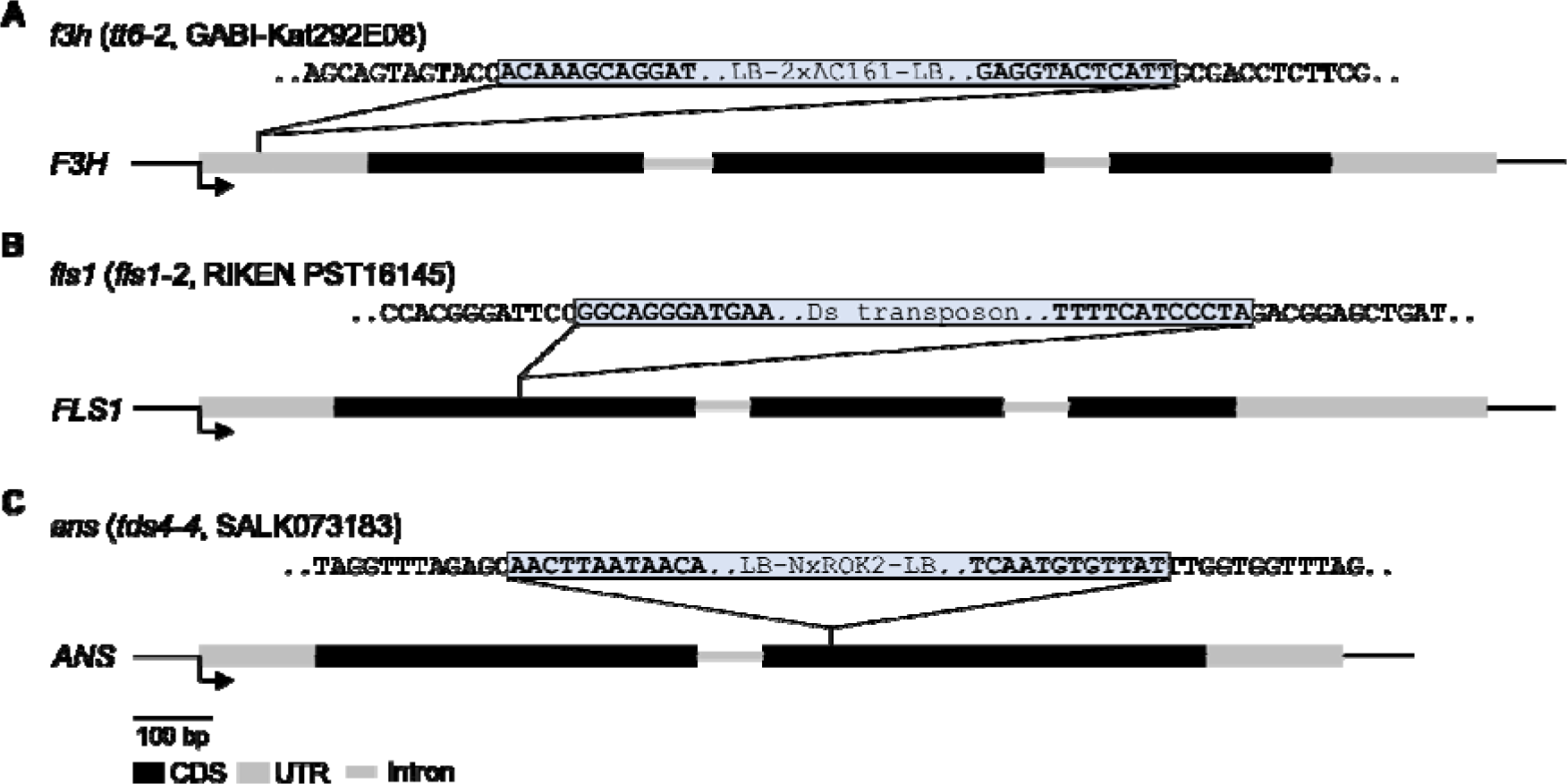
Genomic characterisation of the *f3h*/*fls1*/*ans* mutant. Schematic drawing of the (A) *F3H* (*AT3G51240*), (B) *FLS1* (*AT5G08640*) and (C) *ANS* (*AT4G22880*) genes illustrating the mutations in *f3h* (*tt6-2*), *fls1* (*fls1-2*) and *ans* (*tds4-4*), respectively. Black boxes mark the CDSs and grey boxes indicate UTRs and grey thin lines indicate introns, bent arrows indicate the transcription start. Dashed lines mark the respective T-DNA or transposon insertion. Sequences belonging to the T-DNA or the transposon insertion are shown in a blue box in bold and are flanked by the corresponding genomic sequences. LB, left border.

### Transcriptome analysis of the *f3h*/*fls1*/*ans* mutant

To validate the insertion mutant alleles at the transcriptional level and to examine the transcriptional changes that occur in the *f3h*/*fls1*/*ans* mutant, RNA-Seq data from six-days-old seedlings were analysed. Col-0 and Nö-0 seedlings were used as controls. First, the expression profiles of the respective knockout alleles were analysed in the *f3h*/*fls1*/*ans* mutant (Supplementary Figure S2). For the *ans* and *fls1* mutant alleles, only marginal read coverage was observed in the upstream regions of the T-DNA and transposon insertion positions, respectively. These transcripts, which are greatly truncated compared to their normal lengths, most likely result in non-functional proteins. Surprisingly, the *f3h* mutant allele showed a similar expression profile as the wildtype allele, indicating that the *F3H* locus is still transcribed in the *f3h*/*fls1*/*ans* mutant. However, mapping of the *f3h*/*fls1*/*ans* RNA-Seq reads to the *f3h*/*fls1*/*ans* genome assembly revealed chimeric RNA-Seq reads containing the T-DNA LB sequence and the sequence of the 5’UTR of *f3h*, emphasising that part of the T-DNA sequence is also transcribed (Supplementary Figure S2). Several alternative translation start codons were identified upstream of the original start site of *f3h* (Supplementary Figure S3), which could be used for translation from the mutant *f3h* primary transcript and ultimately lead to a frameshift and/or premature stop codon, resulting in a non-functional protein (refer to the discussion for details).

In total, 1,643 and 1,203 differentially expressed genes (DEGs) were identified when comparing *f3h*/*fls1*/*ans* with Col-0 and Nö-0, respectively (Figure 3, Supplementary File S1). Of the DEGs derived from the comparison between *f3h*/*fls1*/*ans* and Col-0, 768 were up-regulated and 875 were down-regulated, while 818 genes were up-regulated and 385 genes were down-regulated when *f3h*/*fls1*/*ans* was compared to Nö-0. To investigate the transcriptional changes occurring in the *f3h*/*fls1*/*ans* mutant, 86 genes which are differentially expressed between *f3h*/*fls1*/*ans* and Col-0 and between *f3h*/*fls1*/*ans* and Nö-0, but not additionally between Col-0 and Nö-0, were further analysed (Figure 3). After a manual screening (see Methods for details), 35 DEGs remained, which are listed in Table 1. Of these, 28 were up-regulated in the *f3h*/*fls1*/*ans* mutant. The *FLS1* gene was the only known gene related to flavonoid biosynthesis in the list of DEGs.

**Figure 3:**
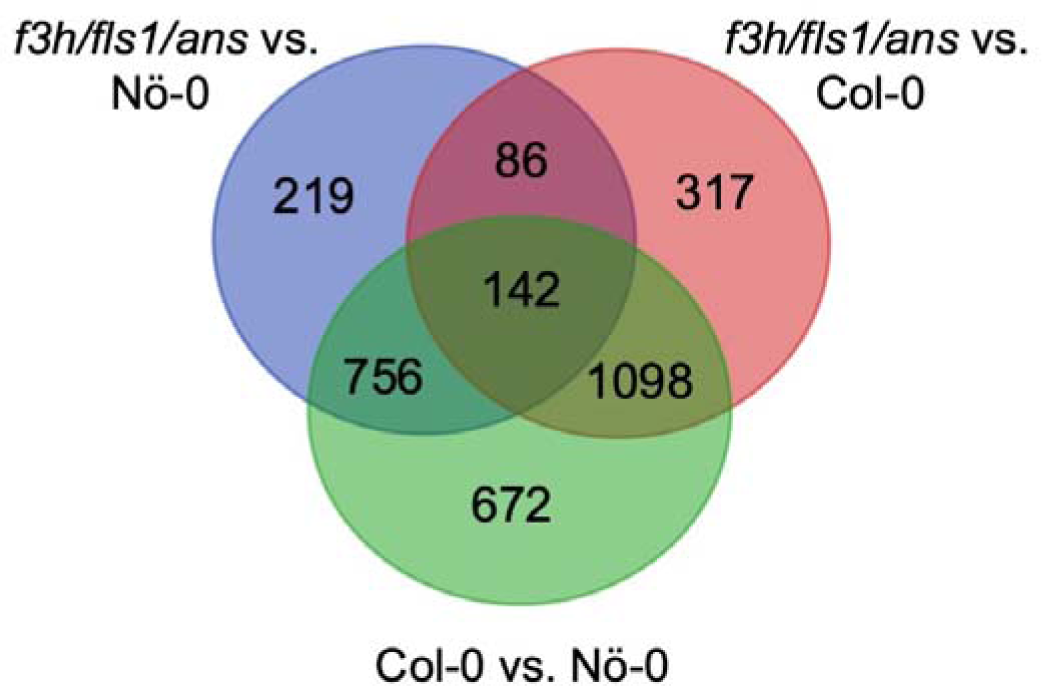
Venn diagram summarising the number of differentially expressed genes between seedlings of *f3h*/*fls1*/*ans* and the wildtypes Col-0 and Nö-0.

**Table 1:** List of differentially expressed genes in *f3h*/*fls1*/*ans* vs. Col-0 and Nö-0. Listed are a subset of differentially expressed genes, that were identified as DEGs in both, the *f3h*/*fls1*/*ans* vs. Col-0 and *f3h*/*fls1*/*ans* vs. Nö-0 comparisons, but notably, they were not differentially expressed between Col-0 and Nö-0. AGI, Arabidopsis gene identifier; l2fc, log_2_ fold change; n.a., no annotation available.

| AGI | l2fc vs. Col-0 | l2fc vs. Nö-0 | name / annotation |
| --- | --- | --- | --- |
| AT2G46765 | 7,25 | 5,74 | hypothetical protein |
| AT5G08640 | -5,54 | -5,71 | FLS1, flavonol synthase 1 |
| AT2G20160 | -4,71 | -5,31 | MEO, E3 ubiquitin ligase SCF complex subunit SKP1/ASK1 family protein |
| AT3G59884 | 3,35 | 1,84 | MIR827A |
| AT2G14610 | 3,31 | 3,64 | PR1 / CAPE9, pathogenesis-related protein 1 |
| AT2G08820 | 2,3 | 1,37 | n.a. |
| AT5G43570 | 2,22 | 1,38 | PR-6 proteinase inhibitor family. |
| AT4G13493 | 2,21 | 1,64 | MIR850A |
| AT4G06075 | 2,15 | 2,20 | MIR5026 precursor |
| AT2G08825 | 2,11 | 1,40 | n.a. |
| AT5G08855 | 2,11 | 1,96 | n.a. |
| AT3G48560 | 2,08 | 2,14 | AHAS / ALS, acetohydroxy acid synthase |
| AT3G46900 | -1,98 | -1,69 | COPT2, Copper transporter 2, copper ion transmembrane transporter |
| AT1G52070 | -1,93 | -1,67 | mannose-binding lectin superfamily protein |
| AT2G03850 | 1,93 | 1,07 | LEA, late embryogenesis abundant protein family protein |
| AT2G11810 | 1,93 | 1,36 | MGD3 / MGDC monogalactosyl diacylglycerol synthase 3 |
| AT2G20723 | 1,84 | 1,79 | snoRNA |
| AT2G20722 | 1,7 | 1,65 | snoRNA |
| AT3G54730 | 1,62 | 3,16 | transcription repressor |
| AT2G20670 | 1,52 | 1,62 | DUF506, putative sugar phosphate exchanger |
| AT3G04210 | 1,36 | 1,59 | TN13, disease resistance protein |
| AT2G44470 | 1,35 | 1,49 | BGLU29, beta glucosidase 29 |
| AT2G02850 | 1,34 | 1,16 | ARPN, plantacyanin, copper ion binding |
| AT1G07135 | 1,29 | 1,23 | glycine-rich protein |
| AT4G39675 | 1,28 | 1,32 | hypothetical protein |
| AT1G60095 | -1,27 | -1,36 | mannose-binding lectin superfamily protein |
| AT3G61930 | 1,26 | 1,17 | hypothetical protein |
| AT2G07698 | -1,23 | -1,91 | ATPase, F1 complex, alpha subunit protein |
| AT2G29330 | 1,23 | 1,54 | TRI, tropinone reductase |
| AT1G36060 | 1,18 | 1,03 | TG / WIND3, translucent green, Integrase-type DNA-binding superfamily protein |
| AT4G18980 | 1,18 | 1,58 | AtS40-3 |
| AT3G44260 | 1,17 | 1,13 | CAF1A, Polynucleotidyl transferase, ribonuclease H-like superfamily protein |
| AT4G28420 | 1,14 | 1,62 | tyrosine transaminase family protein |
| AT3G19430 | -1,12 | -1,90 | LEA protein-like protein, late embryogenesis abundant protein-related |
| AT4G24570 | 1,01 | 1,10 | DIC2, dicarboxylate carrier 2 |

### Phenotypic characterisation of the 2-ODD mutant collection including the novel *f3h*/*fls1*, *f3h*/*ans*, and *f3h*/*fls1*/*ans* mutants

To phenotypically characterise the flavonoid composition of the *f3h*/*fls1*/*ans* mutant compared to the corresponding wildtypes, single and double mutants, the accumulation of flavanones, flavonols, anthocyanins and proanthocyanidins (PA) was analysed (Figure 4, Figure 5). First, the accumulation of flavonols was analysed in norflurazon-bleached seedlings, stained with diphenylboric acid 2-aminoethyl ester (DPBA) (Figure 4A). Both wildtypes, Col-0 and Nö-0, showed an orange colouration indicating the presence of quercetin glycosides. The *chs* mutant, included as a control, is deficient in flavonol glycosides and thus appeared blue due to sinapic derivatives. However, the *ans* mutant displayed an even more intense orange fluorescence than the wildtypes. While the *f3h* mutant showed a red fluorescence, the *fls1* mutant appeared green to light yellow. The *fls1*/*ans* mutant showed a pale yellow colour. The novel *f3h*/*fls1*, *f3h*/*ans*, and *f3h*/*fls1*/*ans* mutants appeared red like the *f3h* mutant.

**Figure 4:**
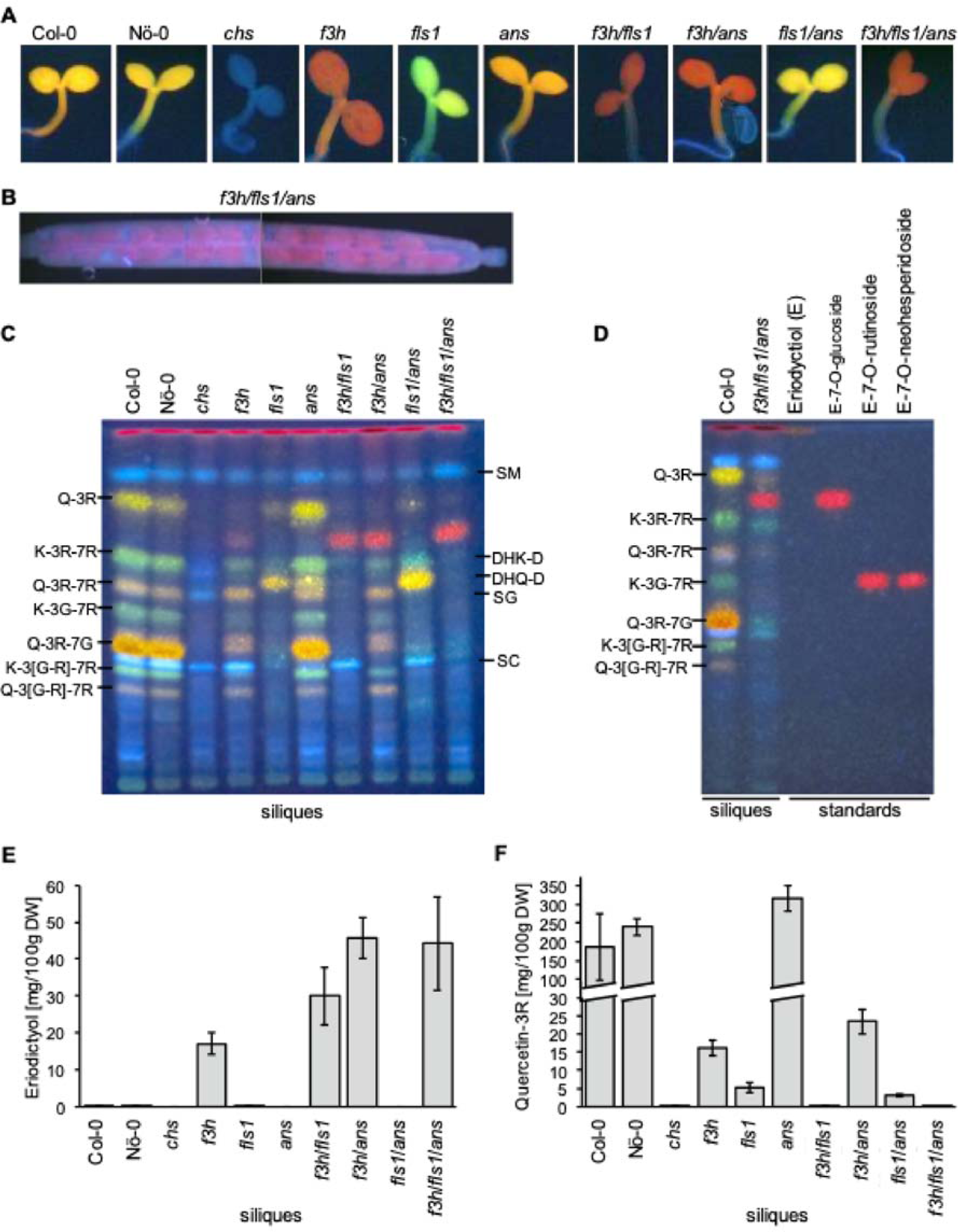
Accumulation of an eriodictyol hexoside in *A. thaliana* 2-ODD mutants carrying the *f3h* allele. (A) Seedlings stained *in situ* with diphenylboric acid 2-aminoethyl ester (DPBA) and visualised under UV light. Quercetin derivatives appear orange, kaempferol derivatives green, and blue corresponds to sinapic acid derivatives. (B) *In situ* DPBA-stained silique of the *f3h*/*fls1*/*ans* mutant visualised under UV light. (C) Accumulation of flavonol- and dihydroflavonol glycosides in siliques analysed by HPTLC and visualised under UV light after treatment with DPBA. The loading to each lane corresponds to methanolic extracts from 3 mg fresh weight of siliques. Kaempferol derivatives are green, quercetin derivatives are orange and sinapic acid derivatives are blue. G, glucose; K, kaempferol; Q, quercetin; R, rhamnose; DHK-D, dihydrokaempferol derivative; DHQ-D, dihydroquercetin derivative; SM, sinapoylmalate; SG, sinapoylglucose; SC, sinapoylcholin. Variations in SG and SC levels are not due to the different genotypes, but to small variations in the developmental stage of the siliques analysed (55). (D) Eriodictyol standards analysed by HPTLC and visualised under UV light after treatment with DPBA. E, eriodictyol. (E) Eriodictyol and (F) quercetin-3-*O*-rhamnoside contents in siliques analysed by UPLC-MS/MS MRM. Error bars indicate the standard deviation of four independent measurements. The original, unprocessed versions of Figures 4C and 4D can be found in Supplementary Figure S13.

**Figure 5:**
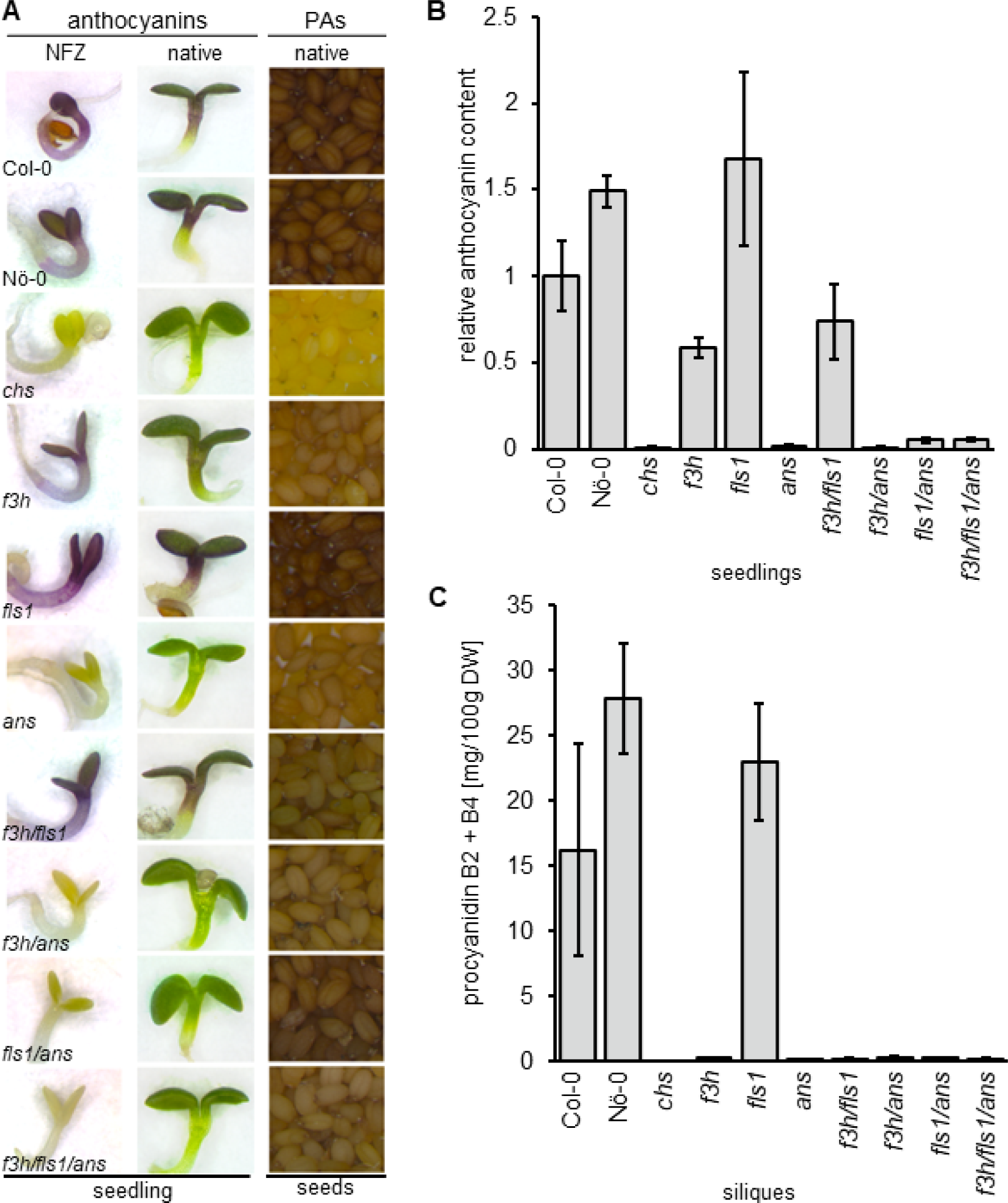
Anthocyanin and proanthocyanidin accumulation in *A. thaliana* 2-ODD mutants. (A) First column: norflurazon (NFZ)-bleached seedlings grown on 4% sucrose for anthocyanin induction. Second column: anthocyanin accumulation in seedlings grown on 0.5 MS plates with 4% sucrose. Third column: native seed colour. (B) Sucrose-induced anthocyanin accumulation in 6-days-old *A. thaliana* seedlings. Three independent biological replicates were measured for each sample. Error bars indicate standard deviation of the mean anthocyanin levels. (C) Procyanidin B2 + B4 content in siliques analysed by UPLC-MS/MS. Error bars indicate the standard deviation of four independent measurements.

HPTLC and targeted UPLC-MS/MS analysis (including MRM quantification) of the flavonoid composition in siliques revealed a wide range of flavonol glycosides in both wildtypes (Figure 4C, Supplementary Table S3). Consistent with DPBA staining, the *chs* mutant was deficient in flavonoids. Accordingly, the level of *p*-coumaric acid was on average 186 µg per g dry weight in *chs* siliques compared to up to 17 µg in all other lines, showing that this flavonoid precursor accumulates in the *chs* mutant as the flavonoid biosynthesis pathway is blocked (Supplementary Table S3). The *f3h*, *ans* and *f3h*/*ans* mutants showed a similar pattern of flavonol glycosides as the wildtypes (Figure 4C). Moreover, the *ans* mutant accumulated the highest amounts of flavonol glycosides with up to 3,549 µg quercetin-3-*O*-rhamnoside (Q-3R) per g dry silique weight, whereas it was <273 µg in the other mutant lines and <2,656 µg in the wildtypes (Supplementary Table S3). Accordingly, the *ans* knockout re-routes flavonoid production to the flavonol biosynthesis branch while the other knockouts cause reduced flavonol levels. The *fls1* mutant accumulated dihydroflavonols and small amounts of Q-3R, whereas Q-3R levels decreased further to an average of 31.3 µg per g dry weight in *fls1*/*ans* siliques while dihydroflavonol levels increased (Figure 4C and F, Supplementary Table S3). The *fls1*/*ans* mutant accumulated the highest dihydroflavonol amounts compared to all lines analysed, with an average dihydroquercetin level of 1,845 µg per g dry silique weight, showing that dihydroflavonols accumulate when a further conversion to flavonols or anthocyanins is not possible. Surprisingly, the *f3h*/*fls1* and *f3h*/*fls1*/*ans* mutants showed marginal and comparable dihydroflavonols levels of up to 55 µg dihydrokaempferol per g dry weight, indicating the presence of an (unknown) 2-ODD enzyme with F3H activity (Supplementary Table S3).

Interestingly, all mutant lines carrying the *f3h* allele accumulated a ‘red (fluorescent) metabolite’, seen between Q-3R and K-3R-7R in HPTLC, indicating that a disruptive *f3h* allele is essential for its production (Figure 4C). These results were in line with the red colouration of *in situ* DPBA-stained seedlings carrying the *f3h* allele (Figure 4A). In contrast to flavonol glycosides and dihydroflavonols, the accumulation of the ‘red metabolite’ was only visible late after DPBA staining. The accumulation of the ‘red metabolite’ was further investigated in DPBA-stained siliques under white and UV light (Supplementary Figure S4). The ‘red metabolite’ visibly accumulated in immature seeds (Figure 4B) of all lines carrying the *f3h* knockout allele and was also observed under white light. The *f3h*/*ans* and *f3h*/*fls1*/*ans* mutants showed the strongest intensity and the ‘red metabolite’ appeared to accumulate during seed maturation (Supplementary Figure S4). In addition, it was detected in young stamens of mutants harbouring the *f3h* allele (Supplementary Figure S5). Interestingly, the eriodictyol content of all analysed lines measured by UPLC-MS/MS were similar to the accumulation pattern of the ‘red metabolite’ observed by HPTLC (Figure 4C and E). In detail, the average level of eriodictyol was <1.4 µg per g dry silique weight in lines with a functional *F3H* allele, whereas the average eriodictyol content of *f3h* knockout lines increased to 170 µg in *f3h*, 300 µg in *f3h*/*fls1,* 457 µg in *f3h*/*ans* and 442 µg in *f3h*/*fls1*/*ans*, indicating that eriodictyol accumulates as a consequence of the *f3h* knockout (Figure 4E). To further characterise the ‘red metabolite’, HPTLC was performed with available eriodictyol glycoside standards together with methanolic extracts of Col-0 and *f3h*/*fls1*/*ans* siliques (Figure 4D). The ‘red metabolite’ had the same retention factor (R_F_) as eriodictyol-7-*O*-glucoside, while the R_F_ values of the eriodictyol disaccharides and the eriodictyol aglycone differed from that of the ‘red metabolite’. Taken together, these results suggest that the metabolite underlying the red colouration of mutants carrying the *f3h* allele is likely to be an eriodictyol hexoside, e.g., eriodictyol-7-*O*-glucoside.

Further chromatographic analysis with ion mobility separation in a high resolution mass spectrometer (UPLC-TWIMS-QToF) on the *f3h*/*fls1*/*ans* flavonoid profile in siliques confirmed the presence of eriodictyol-7-*O*-glucoside compared to a standard, the results show a mass error below 3 ppm and an average of delta collision cross section (CCS) delta of 0.06 % in ESI+ and 2.23 % in ESI- mode (Supplementary Table S4). The average content of eriodictyol-7-*O*-glucoside determined by UPLC-MS/MS (high resolution mass spectrometry) in the *f3h*/*fls1*/*ans* triple mutant is 10.38 µg per g dry weight (Supplementary Table S5). Interestingly, the profile in *f3h*/*fls1*/*ans* is more complex (Supplementary Figures S6-S8). The annotation of the eriodictyol derivates profile are based on HDMS^E^ data, fragmentation patterns by UPLC-MS/MS and isotopic distribution. The UPLC-TWIMS-QToF HDMS^E^ profile obtained in ESI+ and ESI- mode and subsequently UPLC-MS/MS analysis revealed the presence of another eriodictyol hexoside, four di-hexosides, two tri-hexosides and two rutinosides or neohesperidosides, respectively (Supplementary Tables S4 and S6). In ESI- mode the main loss by UPLC-MS/MS is 162 Da (Supplementary Figure S8), corresponding to the loss of a hexoside, the fragmentation has the presence of *m/z* 287, *m/z* 151 and *m/z*135 (Supplementary Figure S8), indicating a core with molecular formula C_15_H_12_O_6_. Eriodictyol and dihydrokaempferol have the same precursor, naringenin (Figure 1), and the same molecular formula. Therefore, a comparison of fragmentation patterns, under the same methodology, of eriodictyol and dihydrokaempferol standards by UPLC-MS/MS and UPLC-HDMS^E^ low and high energy is required to improve the metabolite annotation. As is shown in Supplementary Figures S9-S11, the absence of *m/z* 177 in all compounds of interest, including the eriodictyol-7-*O*-glucoside, indicates an eriodictyol core. Two compounds, tentatively annotated as eriodictyol rutinoside or neohesperidoside have two losses of 162 Da and 308 Da in ESI- and 308 Da in ESI+ mode (Supplementary Figure S8, Supplementary Tables S4 and S6). The presence of eriodictyol-7-*O*-rutinoside is excluded by the mismatch in retention time and CCS values in the triple mutant by HDMS^e^ (Supplementary Table S4) and is in agreement with the results shown in Figure 4D. Moreover, the dihydroquercetin profile of the *f3h*/*fls1*/*ans* triple mutant compared to Col-0 is interesting, as it contains three different hexosides (Supplementary Table S4). These results shed light on the accumulation of the eriodictyol and dihydroquercetin aglycons and their subsequent modifications in the *f3h*/*fls1*/*ans* mutant.

Besides flavanones and flavonols, other downstream (end-)products of the flavonoid biosynthesis were analysed including anthocyanins and PAs. The accumulation of anthocyanins was analysed in sucrose-induced, norflurazon-bleached seedlings (Figure 5A). While both wildtypes, *f3h*, *fls1*, and *f3h*/*fls1* mutants show anthocyanin accumulation, *chs*, *ans*, *f3h*/*ans*, *fls1*/*ans*, and *f3h*/*fls1*/*ans* mutants do not. These results are consistent with the analysis in sucrose-induced, native seedlings and were validated by photometric quantification of anthocyanins in seedlings (Figure 5A and B).

Seed pigmentation is influenced by the levels of PA in the seed coat, with a darker seed colour corresponding to higher PAs levels. Analysis of the native seed colour of the wildtypes revealed characteristic dark brown colour due to the presence of large amounts of PAs, while the *chs* mutant showed a pale yellow seed indicative of the well-known PA deficiency and transparent testa phenotype (Figure 5A). Most of the *2-ODD* mutants showed an intermediate seed colour phenotype. However, the *fls1* mutant showed a similar brown seed colour as the wildtypes. In accordance, UPLC-MS/MS analysis revealed that epicatechin levels of Col-0, Nö-0, and *fls1* averaged between 799 µg and 1,108 µg per g dry silique weight, while all other lines showed average levels <7 µg (Supplementary Table S3). Similar observations were made for procyanidins (Figure 5D, Supplementary Table S3). Thus, PAs and their precursor epicatechin are only produced in considerable amounts when *CHS, F3H*, and *ANS* are intact.

## Discussion

The three major flavonoid 2-ODD enzymes, F3H, FLS and ANS, are closely related and therefore often have overlapping catalytic activities. We have generated an *A. thaliana f3h*/*fls1*/*ans* mutant and analysed genomic, transcriptomic and phenotypic changes in order to increase the knowledge about flavonoid biosynthesis related 2-ODDs in *A. thaliana*.

### Genome sequence of the *f3h*/*fls1*/*ans* triple mutant

To characterise the *f3h*/*fls1*/*ans* triple mutant at the genomic level, a *de novo* Oxford Nanopore Technology (ONT) genome assembly was generated. In-depth analysis of the respective *2-ODD* mutant loci revealed the expected insertion positions per mutant allele (49, 51, 56). To our knowledge, these mutant alleles have never been characterised using long-read technology. Thus, in this study, the insertion of repeated T-DNAs and their BVB sequences at the *ans* (*tds4-4*) locus were identified. Moreover, the ONT assembly revealed that two T-DNAs inserted head-to-head in the 5’UTR of *f3h*. Such complex T-DNA insertion events, involving arrays of multiple T-DNAs as well as BVB sequences, have been reported before to be more the rule than the exception for *A. thaliana* T-DNA insertion mutants (57–60).

### Transcriptome analysis of the *f3h*/*fls1*/*ans* triple mutant

RNA-Seq analysis in seedlings revealed that *FLS1* is the only differentially expressed 2-ODD between the *f3h*/*fls1*/*ans* mutant and the wildtypes and not additionally differentially expressed between the wildtypes Col-0 and Nö-0. *ANS* is downregulated in the triple mutant compared to the two wildtypes, but additionally also downregulated in Nö-0 compared to Col-0. *F3H* was not identified as differentially expressed gene (DEGs) due to remaining RNA-Seq read coverage at the *f3h* loci downstream of the T-DNA insertion site (Supplementary Figure S2). For *ANS*, which comprises two exons, read coverage of the most of the first exon was observed upstream of the complex T-DNA insertion array. The resulting transcript is severely truncated and, as previously confirmed by the phenotype of the homozygous mutant (51, 52), results in a non-functional protein.

Interestingly, the expression level of the *f3h* locus was similar to that of the wildtype alleles and covered the entire *F3H* exons. Comparison of the RNA-Seq data to the *f3h*/*fls1*/*ans* genome assembly identified several chimeric RNA-Seq reads spanning part of the T-DNA and the 5’UTR of *f3h*, emphasising that part of the T-DNA::genome junction is transcribed. Transcription is likely driven by the 2’promoter of the T-DNA, located 723 bp upstream of the 5’UTR of the *f3h* locus (Supplementary Figure S3). A similar observation was made in a *phytochrome C* mutant by *Monte et al.* (61). They analysed a T-DNA insertion line that showed *PHYC* transcript levels similar to the wildtype. However, protein analysis revealed, that the plants did not accumulate the corresponding protein. Similar limitations are conceivable in our study. The 11,393 bp T-DNA insertion could either lead to an extended and, in consequence, non-functional protein or one of the start codons located on the T-DNA could be used as an alternative start codon, ultimately resulting in a frameshift and/or premature stop codon (Supplementary Figure S3). Taken together, the omics data and the very distinct phenotype of the *f3h*/*fls1*/*ans* mutant strongly suggest a non-functional *f3h* allele, in line with previous results (17, 56). Furthermore, genomic and transcriptomic data of the *ans* and *fls1* loci showed that these loci are *null*-alleles.

Apart from the *FLS1* gene, no other genes encoding enzymes, regulators or transporters of flavonoid biosynthesis were identified as DEGs between the *f3h*/*fls1*/*ans* mutant and the wildtypes. Nevertheless, several identified DEGs are noteworthy. For example, *miR827A* is upregulated in *f3h*/*fls1*/*ans*, compared to the wildtypes. *miR827* expression is known to be sucrose-induced and it targets ethylene response factors (ERFs) to regulate the expression of *SUCROSE SYNTHASE* which has been proposed as a regulator of anthocyanin biosynthetic genes (62). Accordingly, increased *miR827A* expression could be a means of the *f3h*/*fls1*/*ans* mutant to counteract anthocyanin deficiency. Of course, this requires an unknown feedback regulation circuit. *TROPINONE REDUCTASE* (*TRI*) transcript abundance is increased in *f3h*/*fls1*/*ans* seedlings. Moreover, based on publicly available RNA-Seq datasets deposited in of the European Bioinformatics Institute (EBI) Expression Atlas, *TRI* is expressed in Col-0 siliques (63, 64). TRI has previously been reported to reduce the flavanone backbone (65). Since the flavonol- and anthocyanin branches of flavonoid biosynthesis are blocked in the *f3h*/*fls1*/*ans* mutant, flavanones like eriodictyol accumulate in *f3h*/*fls1*/*ans* and could be reduced by TRI activity. *BETA GLUCOSIDASE29* (*BGLU29*) is upregulated in *f3h*/*fls1*/*ans* compared to the wildtypes. BGLU29 belongs to the family of beta-glucosidase enzymes (66). Although some BGLUs have been shown to be functional glucosyltransferases involved in the glycosylation of flavonoids (BGLU10 (67) and BGLU6 (68)), it is very unlikely that BGLU29 has glycosyltransferase activity, as it is most closely related to BGLU28 - BGLU32, all of which having glycoside hydrolase activity (69). BGLU29 may be involved in the hydrolysis of flavonoid glycosides. In addition, the expression of *ACETOHYDROXY ACID SYNTHASE* (*AHAS*), which is involved in the biosynthesis of valine, leucine and isoleucine from pyruvate (70), is increased in the *f3h*/*fls1*/*ans* mutant. Since flavonoids are derived from phenylalanine, increased *AHAS* expression could hint to an accumulation of phenylalanine in *f3h*/*fls1*/*ans* and an effort by the plant to produce other amino acids instead.

### Phenotype

#### *In planta* multifunctionality of flavonoid 2-ODDs

Previous studies revealed a “leaky” phenotype of the *2-ODD* single mutants and the *fls1*/*ans* double mutant, which was attributed to partial compensation of other 2-ODD dioxygenases, e.g., the bi- and multifunctionality of FLS and ANS, respectively (21, 27, 49). Owens *et al.* (27) showed, that FLS1 not only accepts dihydroflavonols as substrate, but can also convert naringenin to dihydrokaempferol *in planta*, thus exhibiting F3H side activity. ANS has been shown to produce dihydroquercetin and quercetin from naringenin *in vitro*, indicating its F3H and FLS side activity (40, 71, 72). Furthermore, the accumulation of flavonol glycosides in the *fls1* mutant and further flavonol reduction in the *fls1*/*ans* mutant showed that ANS also has an FLS activity *in planta* (49). This knowledge is consistent with our analyses and the novel *f3h*/*ans*, *f3h*/*fls1* and *f3h*/*fls1*/*ans* mutants allowed an in-depth analysis of the *in planta* multifunctionality of F3H, FLS, and ANS. The flavonol glycoside patterns of the *f3h*, *ans* and the novel *f3h*/*ans* mutants were like the ones of the wildtypes confirming the significant *in planta* F3H activity of FLS1.

The *fls1*/*ans* mutant still accumulates flavonol glycosides, suggesting that F3H has additional *in planta* FLS activity. Alternatively, FLS3 may have residual *in planta* FLS activity, as FLS *in vivo* activity of FLS3 has been shown in yeast (73). As all mutant lines with a non-functional ANS contain no anthocyanins, F3H and FLS do not seem to have any ANS activity. Finally, the marginal and comparable amounts of dihydroflavonol of up to 55 µg dihydrokaempferol per g dry silique weight identified in the *f3h*/*fls1* and *f3h*/*fls1*/*ans* mutants indicate that ANS does not appear to have significant F3H activity *in planta*. However, the identification of residual dihydroflavonols in the *f3h*/*fls1*/*ans* mutant was rather surprising since the two major flavonoid 2-ODD responsible for dihydroflavonol accumulation, F3H and FLS1, were knocked-out in this mutant.

Although the biological significance of these trace amounts of dihydroflavonols and the enzyme(s) leading to their production is questionable, it raises the question of which enzyme(s) could be involved in their biosynthesis. The dihydroflavonols could be produced by an (unknown) 2-ODD with F3H activity or F3H side activities of one of the FLS paralogs. While *AthFLS2*, *AthFLS4*, *AthFLS5* and *AthFLS6* were identified as pseudogenes or encoding non-functional proteins, minor FLS activity of *Ath*FLS3 was reported under prolonged assay conditions in yeast (27, 73). Although FLS3 showed no F3H or FLS activity in *Escherichia coli*, *in planta* F3H activity was identified for FLS3 homologs from banana and rapeseed, the latter sharing 67 % amino acid sequence identity with *Ath*FLS3 (42, 46). To investigate a potential *in planta* F3H and FLS activity of *Ath*FLS3, an *fls3* knockout allele could be introduced in the *f3h*/*fls1*/*ans* mutant to generate a *fls3*/*f3h*/*fls1*/*ans* mutant. A reduction of dihydroflavonols and flavonols in the *fls3*/*f3h*/*fls1*/*ans* mutant compared to the *f3h*/*fls1*/*ans* mutant would provide evidence for an F3H and FLS activity of FLS3 *in planta*.

Alternatively, the production of residual dihydroflavonols in the *f3h*/*fls1*/*ans* mutant may involve FNSI, another flavonoid 2-ODD closely related to F3H. Bifunctional enzymes with FNSI and F3H activity have been identified in gymnosperms and mosses (74). FNSI enzymes were thought to be restricted to the Apiaceae family (75) and thus absent from *A. thaliana,* until the identification of a DOWNY MILDEW RESISTANT6 (DMR6/FNSI) homolog with FNSI activity, alongside reports of functional FNSIs from liverworts, *Oryza sativa*, and *Zea mays* (26, 74, 76, 77). However, *DMR6*/*FNSI* is primarily expressed in leaves and not in siliques (26), where the residual amounts of dihydroflavonols were detected. Moreover, the endogenous activity of DMR6/FNSI is not sufficient to compensate the dihydroflavonol deficient phenotype of the *f3h*/*fls1* or *f3h*/*fls1*/*ans* mutants, indicating that DMR6/FNSI does not have substantial F3H activity *in planta*. Taken together, these findings suggest that if DMR6/FNSI harbours F3H side activity, it is likely to play a minor role, if any, in the production of the residual dihydroflavonols detected in *f3h*/*fls1*/*ans* siliques.

In summary, the established 2-ODD mutant collection could be used in the future to analyse not only potential multifunctionality of DMR6/FNSI, but rather 2-ODDs in general. This is of particular importance for the FNSI family, as it is no longer confined to the Apiaceae family, and knowledge on FNSI multifunctionality is rare, especially in angiosperms (74, 78). As mentioned above, flavonoid 2-ODDs show broad substrate specificities *in vitro,* which do not necessarily correspond to product accumulation *in vivo* (21). This emphasises the importance of the established mutant collection for the *in planta* characterisation of flavonoid 2-ODD multifunctionality, especially for studies characterising their functionalities solely *in vitro* (79, 80). For example, *in planta* complementation experiments using the novel mutant collection could validate the FLS side activity found *in vitro* and the absence of F3H side activity of *Vitis vinifera*’s ANS (80).

#### Flavonoid accumulation in the absence of the three major flavonoid 2-ODDs

The mutant collection was harnessed to investigate changes in the metabolic flux associated with the knockout of the three major flavonoid 2-ODDs F3H, FLS1, and ANS. Interestingly, all mutant lines carrying the *f3h* allele accumulate a metabolite in seedlings, immature seeds, young stamens and siliques, which appears ‘red’ after DPBA-staining in visible light and under UV light. This ‘red (fluorescent) metabolite’ is absent in the wildtypes and is most clearly seen in an HPTLC of methanolic extracts from siliques (Figure 4B). The characteristic red fluorescence of the *f3h* seedlings after DPBA staining has previously been observed and suggested to be derived from an orthodihydroxy- or orthotrihydroxy flavonoid (81). In addition, 3’hydroxylation of the B ring was suggested to be required for the red fluorescence upon staining, as DPBA-stained *f3h*/*f3’h* and *chs*/*f3h* mutants showed a light blue to pale fluorescence (81). Owens *et al.* (21) suggested that F3’H might convert the accumulating naringenin to eriodictyol which could be further reduced to a 3-desoxy flavonoid by the FNR activity of native DFR (21). Wollenweber (82) postulated that the metabolite underlying the red spot on DPBA-stained TLC plates is an eriodictyol derivative. We show that high levels of eriodictyol accumulate in siliques of lines carrying an *f3h* knockout allele. This is in accordance with the high expression of *F3’H* in siliques compared to flowers, leaves, stems, and roots (22). In addition, we also observed that *F3’H* is expressed in *f3h*/*fls1*/*ans* seedlings (Supplementary Figure S12), which showed the characteristic red colouration after *in situ* DPBA-staining. While HPTLC-based flavonoid glycoside analysis showed that the ‘red metabolite’ is most likely an eriodictyol hexoside, the results by UPLC-MS/MS and UPLC-TWIMS-QToF analyses revealed a profile with various metabolites annotated as eriodictyol hexosides, including eriodictyol-7-*O*-glucoside, and additional possible eriodictyols with neohesperidoside or rutinoside derivatisation.

We therefore postulate that F3’H converts naringenin to eriodictyol, which may then be further modified by glycosylation via glycosyltransferases. For example, the flavonoid-7-*O*-glycosyltransferase UGT73B1 has been shown to use eriodictyol as a preferred substrate for the catalysis to eriodictyol-7-*O*-glycoside (35). Considering the previously presented DEGs, the tropinone reductase (TRI) may be involved in eriodictyol reduction, as a previous study showed that TRI can reduce flavanones *in vitro* (83). In the future, knockout alleles of the identified candidate genes could be introduced in the *f3h*/*fls1*/*ans* mutant to analyse their role in the eriodictyol hexoside biosynthesis and degradation.

## Conclusions

We present a detailed characterisation of a novel *A. thaliana f3h*/*fls1*/*ans* mutant and the corresponding single and double mutants. We have used genomic, transcriptomic, and metabolite data to extend the knowledge about flavonoid 2-ODDs in *A. thaliana.* The set of *f3h, fls1* and *ans* single, double and triple mutants will enable further work on flavonoid 2-ODDs and their multifunctionalities in *A. thaliana*, as these are often limited to *in vitro* studies or enzyme assays in *E. coli* or yeast (45, 79, 84). The *f3h*/*fls1*/*ans* mutant will also facilitate the characterisation of multifunctional 2-ODDs from other species *in planta*. Finally, we have shown that eriodictyol and several eriodictyol derivatives accumulate in the absence of a functional F3H *in planta* with eriodictyol-7-*O*-glucoside being one of the predominant eriodictyol derivatives. (Homo-)eriodictyols are bitter-masking flavanones with no additional strong taste or flavour and are thus promising flavour modifiers for food applications and pharmaceuticals (85).

## Methods

### Plant material

Columbia-0 (Col-0, NASC ID N1092) and Nössen-0 (Nö-0, NASC ID N3081) accessions were used as wildtype controls. A *chs* mutant (*tt4-11*, SALK_020583, Col-0 background, T-DNA from pROK2 plasmid (18, 86)) was used as a flavonoid deficient control. The *A. thaliana* mutants *f3h* (*tt6-2*, GK-292E08, Col-0 background, T-DNA from pAC161 plasmid (17)), *fls1* (*fls1-2,* RIKEN_PST16145, Nö-0 background, Ds transposon (16, 49)), *ans* (*tds4-4*, SALK_073183, Col-0 background (51, 52)) and an *fls1*/*ans* double mutant (49) were used for crosses and as controls.

### Crossing and genotyping of mutants

To generate the novel *f3h*/*fls1*, *f3h*/*ans*, and *f3h*/*fls1*/*ans* mutants, the *fls1*/*ans* double mutant was crossed with the *f3h* mutant. Individuals of the F2 progeny were analysed by PCR genotyping using GoTaq2 DNA polymerase (Promega) according to standard protocols. Sequences of primers used for genotyping are provided in Supplementary Table S2. Plants homozygous for the respective mutant alleles were used for further analyses.

### DNA extraction, library construction, and sequencing

ONT was used to generate long-reads of the *f3h*/*fls1*/*ans* mutant. Genomic DNA was isolated from rosette leaves using the NucleoBond HMW DNA Kit (Macherey-Nagel, Düren, Germany) according to the manufacturer’s instructions. DNA quality was assessed via PicoGreen (Quant-iT) measurement on a FLUOstar OPTIMA plate reader (BMG labtech), and pulsed-field gel electrophoresis. The Short Read Eliminator (SRE) Kit (Circulomics, Baltimore, MD, USA) was used to enrich long DNA fragments and results were validated by PicoGreen measurements. 1.2 µg of high molecular weight DNA was used for library preparation according to the SQK-LSK109 protocol (Oxford Nanopore Technologies (ONT), Oxford, UK). Sequencing was performed on a R9.4.1 flow cell on a GridION. The flow cell was treated with nuclease flush to load another library twice. Basecalling was performed using Guppy v.4.2.3 (https://community.nanoporetech.com).

### *De novo* ONT genome assembly and read mapping

A *de novo* ONT genome assembly of the *f3h*/*fls1*/*ans* mutant was constructed with Canu v2.1.1 (87). No polishing of the assembly was performed, as the assembly was only used to analyse the respective mutant loci. Contigs <100 kb were removed, followed by the calculation of assembly statistics as previously described (88). Assembly quality was assessed via BUSCO v3.0.2 using the embryophyta_odb9 lineage dataset (89). For manual curation of loci of interest reads were mapped to the *A. thaliana* TAIR10 reference genome sequence (90) using minimap2 (91). Minimap2 was run in ONT mode (-ax map-ont) and no secondary alignments were allowed (--secondary=no). BAM files were sorted and indexed using samtools (92). Manual curation of regions of interest was performed by using Integrative Genomics Viewer (93) and BLAST (94). The assembly data was screened for the presence of additional T-DNAs using loreta (58).

### RNA-Seq analysis: Ribonucleic acid extraction, library construction, and sequencing

RNA was isolated from 6-days-old seedlings using the NucleoSpin^®^ RNA Plant Kit (Macherey-Nagel, Düren, Germany) according to the manufacturer’s instructions. Four biological replicates were used for each genotype (Col-0, Nö-0, and *f3h*/*fls1*/*ans*). The RNA quality was validated using a 2100 Bioanalyzer system (Agilent, Santa Clara, CA, USA). Sequencing libraries were constructed from 1 µg of total RNA, following the TrueSeq v2 protocol. Single end sequencing of 84 nt was performed on an Illumina NextSeq500 at the Sequencing Core Facility of the Center for Biotechnology (CeBiTec) at Bielefeld University.

### RNA-Seq mapping and count table generation

After assessing read quality using FastQC (95), reads were mapped to the *A. thaliana* Col-0 TAIR10 reference genome sequence (96) using STAR v.2.7.1a (97). STAR was run in Basic mode allowing a maximum of 5 % mismatches per read length and using a minimum of 90 % matches per read length. Mapping statistics were calculated based on STAR.log files using a customised python script (available online via https://github.com/hschilbert/3xODD/blob/main/parse_STAR_log_file_create_mapping_statis tic.py). We used featureCounts v2.0.1 (98) with the -O --fraction parameters to generate count tables based on the *A. thaliana* araport11 gff3 file (99) in order to assign a fractional count of 1/y from a reads, where y is the total number of meta-features/features overlapping with the reads.

### Analysis of differentially expressed genes

To identify differentially expressed genes (DEGs) between Col-0, Nö-0 and the *f3h*/*fls1*/*ans* mutant in young seedlings, DESeq2 (100) was used to identify DEGs with a log_2_ fold change >1 and adjusted p-value <0.05 were selected for downstream analysis. Selected DEGs (86 DEGs derived from the comparison of *f3h*/*fls1*/*ans* mutant to both wildtypes) were manually curated by in-depth analysis of DEG’s RNA-Seq read coverage against the TAIR10 reference genome sequence to exclude DEGs arising from mapping artefacts, e.g., genes with partial read coverage. The Venn diagram was generated using https://bioinformatics.psb.ugent.be/webtools/Venn/.

### Flavonoid staining in *A. thaliana*

Visualisation of DPBA/Triton X-100-stained flavonoids in norflurazon-bleached *A. thaliana* seedlings was performed according to Stracke *et al.* (101) using epifluorescence microscopy. Visualisation of flavonoids in ethanol-bleached siliques of adult *A. thaliana* plants was performed according to Stracke *et al.* (102).

### High performance thin layer chromatography (HPTLC) of flavonols and dihydroflavonols

Plants for flavonols extraction were grown in a greenhouse with 16 h of light illumination per day at 22 °C. Extraction and analysis of dihydroflavonol and flavonol glycoside compounds were performed as described in Stracke *et al.* (49). A total of 9-10 siliques of similar length were harvested from each genotype, yielding 22-56 mg fresh weight material. Samples were homogenised in 80 % methanol, incubated at 70 °C for 15 min and centrifuged at 14,000 x g for 10 min. After vacuum-drying of the resulting supernatants, the dried pellets were redissolved in 80 % methanol (1µL/mg) and 3 µL were analysed by HPTLC on a Silica Gel 60 plate (Merck, Darmstadt, Germany). The mobile phase used was ethyl acetate, formic acid, acetic acid and water (100:26:12:12). Dihydroflavonols and flavonol glycosides were detected by DPBA staining and UV light.

### (Poly)phenol extraction and targeted Ultra Performance Liquid Chromatography – Multiple Reaction Monitoring Mass Spectrometry (UPLC-MS/MS MRM) analysis

Samples of plant material were taken directly in the laboratory, lyophilized, and stored at room temperature until used for further analysis. 100 mg of ground tissue were mixed with 1 mL of 80 % methanol for extraction of (poly)phenolic compounds. Samples and solvent were mixed on an orbital shaker for 1 h at room temperature, sonicated for 30 min and the extraction continued overnight at 4 °C in the dark. Each extract was filtered through a MILLEX 13-0.22 µm PTFE membrane into glass vials and analysed as follows.

Sample extracts were injected immediately after filtration. Targeted UPLC-MS/MS with MRM quantification analysis was performed on a Waters Acquity UPLC system (Milford, MA, USA) consisting of a binary pump, an online vacuum degasser, an autosampler and a column compartment. Separation of the phenolic compounds was conducted on a Waters Acquity HSS T3 column 1.8 μm, 100 mm × 2.1 mm, kept at 40 °C. The analysis of phenolic compounds was performed according to the method described by Vrhovsek *et al.* (103) using water with 0.1 % formic acid and acetonitrile with 0.1 % formic acid as mobile phases for the gradient.

Mass spectrometry detection was performed on a Waters Xevo TQMS instrument equipped with an electrospray ionisation (ESI) source. The capillary voltage was 3.5 kV in positive mode and −2.5 kV in negative mode; the source was maintained at 150 °C; the desolvation temperature was 500 °C; cone gas flow was 50 L/h; and desolvation gas flow was 800 L/h. Unit resolution was applied to each quadrupole. Prior to UPLC-MS/MS MRM analysis, infusion of each individual standard was performed to optimize the MRM conditions for quantification. Data processing was performed using the Mass Lynx TargetLynx Application Manager (Waters).

### Ultra-Performance Liquid Chromatography - Traveling Wave Ion Mobility-Mass Spectrometry - Quadrupole Time of Flight (UPLC-TWIMS-QToF) Mass Spectrometry and tandem mass spectrometry (UPLC-MS/MS) analysis

The metabolite profile of Col-0 and *f3h*/*fls1*/*ans* biological replicate sample extracts were acquired in an UPLC Acquity I system (Waters Corporation) coupled to a high-resolution mass spectrometer HDMS Synapt XS (Waters Corporation). The chromatographic methodology was performed according to Arapitsas *et al.* (104) with some modifications. The chromatographic separation was carried out in an Acquity Premier HSS T3 1.8 µm, 2.1 x 100 mm column (Waters) at 40 °C. The mobile phases A and B consisted of water with 0.1 % of formic acid and methanol with 0.1 % formic acid, respectively, pumped with a flow rate of 0.3 mL/min. High definition MS^e^ (HDMS^e^) data acquisition with a scan time of 0.2 s was performed in continuum mode with a mass range of 50-2000 Da in resolution mode (≥ 21500 FWHM at *m/z* 785), for both ionisation modes (ESI+ and ESI-). The source temperature was settled at 120 °C, the desolvation temperature at 450 °C, the nebuliser gas volume flow was 50 L/h and the desolvation gas volume flow rate was 800 L/h. For ESI-acquisition mode, the capillary voltage was 2.0 kV, and sampling cone was 30 V, whereas for ESI+ mode, the capillary voltage was 3.0 kV, and the sampling cone was 40 V. A collision energy ramp of 20-40 V was applied in the transfer cell for high energy. For the ion mobility separation, the helium cell flow rate was 180 mL/min and for nitrogen it was 90 mL/min. The wave velocity was set to 650 m/s and the transfer wave velocity to 141 m/s. The calibration was performed using the Major Mix IMS/ToF Calibration kit solution (Waters Corporation). A leucine enkephalin solution (200 pg/µL) was continuously infused as lock mass at flow rate of 10 µL/min during the whole analysis. MassLynx v.4.2 and UNIFI software (Waters Corporation) were used for data acquisition and processing. The collision cross sections (CCS) obtained were compared with an *in-house* standard library.

A target UPLC-MSMS analysis was performed with the same source conditions in resolution mode in a range of 50-2000 Da, with a scan time of 1 s. The collision energy was applied as a ramp in the trap from 10-50 V for ESI- and 10-40 V for ESI+.

### Determination of anthocyanin accumulation

Anthocyanin accumulation was visualised in 6-days-old seedlings grown on filter paper soaked with 3 ppm of the bleaching herbicide norflurazon supplemented with 4 % sucrose to induce anthocyanin accumulation. In addition, anthocyanin content was quantified photometrically in 6-days-old seedlings grown on 0.5 MS plates with 4 % sucrose as described by Mehrtens *et al.* (105). Three independent biological replicates were measured for each sample.

## Declarations

### Ethics approval and consent to participate

Not applicable.

### Consent for publication

Not applicable.

### Availability of data and materials

All sequence data generated in this study are available under the ENA/NCBI Bioproject ID PRJEB60353. In detail, the RNA-Seq data of *A. thaliana* Col-0, Nö-0, and the *f3h*/*fls1*/*ans* mutant can be found with under the ENA/NCBI IDs: ERS14716218 - ERS14716221, ERS14716222 - ERS14716225 and ERS14716226 - ERS14716229, respectively. The genomic ONT data can be accessed using the ENA/NCBI IDs: ERS14716230 (FASTQ) and ERS14716231 (FAST5). The *A. thaliana f3h*/*fls1*/*ans* mutant ONT assembly can be accessed at the DOI: 10.4119/unibi/2979807.

### Cometing interests

The authors declare that they have no competing interest.

### Authors’ contributions

MB, HMS, RS and BW conceived and designed the research. MB, HMS, RS and SM carried out experiments and analysed the data. VS performed high resolution mass spectrometry experiments and interpreted the data. AA performed the metabolite quantification methodology. RS and BW supervised the project. MB and HMS drafted the manuscript. RS, SM and BW revised the manuscript.

## Acknowledgments

We thank Andrea Voigt, Melanie Kuhlmann and Nicoletta Schmidt for excellent technical assistance and Martin Sagasser for fruitful discussions. Moreover, we thank Nils Kleinbölting for his support regarding *loreta*. We thank the Center for Biotechnology (CeBiTec) at Bielefeld University for providing an environment to perform the computational analyses.

