## Supplementary Figure for "Generation and characterisation of an *Arabidopsis thaliana f3h*/*fls1*/*ans* triple mutant that accumulates eriodictyol derivatives"

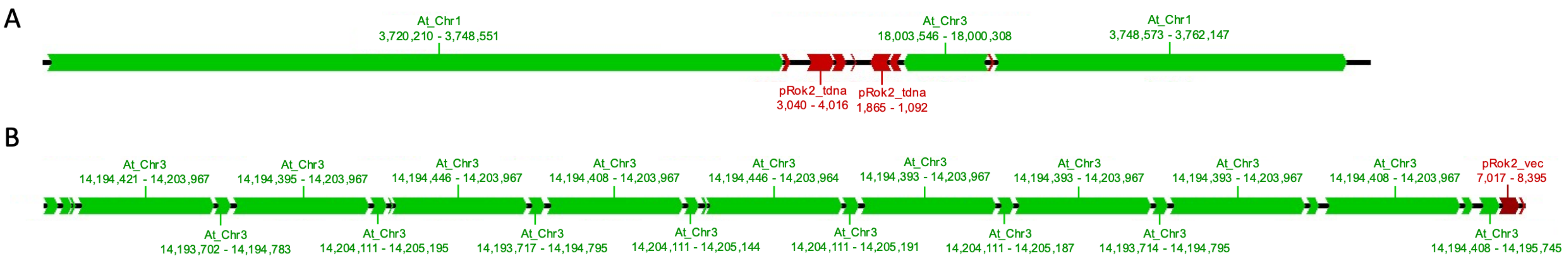


**Supplementary Figure S1: Additional T-DNA insertions in the *f3h*/*fls1*/*ans* mutant identified by loreta.** Schematic drawing of two additional T-DNA insertion events in the *f3h*/*fls1*/*ans* mutant. (A) The first insertion on chromosome one is located between 3,748,551 – 3,748,573 and carries a 3,238 bp inverted fragment of chromosome three consisting of the sequence from 18,003,546 – 18,000,308 bp. Labelling of fragments smaller than 500 bp was omitted for clarity. (B) The second insertion is followed by a large tandem repeat of the chromosome three fragment ranging from ~14,194 – 14,204 kbp. Labelling of fragments smaller than 1,000 bp was omitted for clarity. Green boxes indicate genomic fragments matching the TAIR10 reference genome sequence and red boxes indicate the presence of T-DNA (ending with "_tdna") or binary vector sequences (ending with "_vec"). The arrow shape marks the orientation of the fragments.

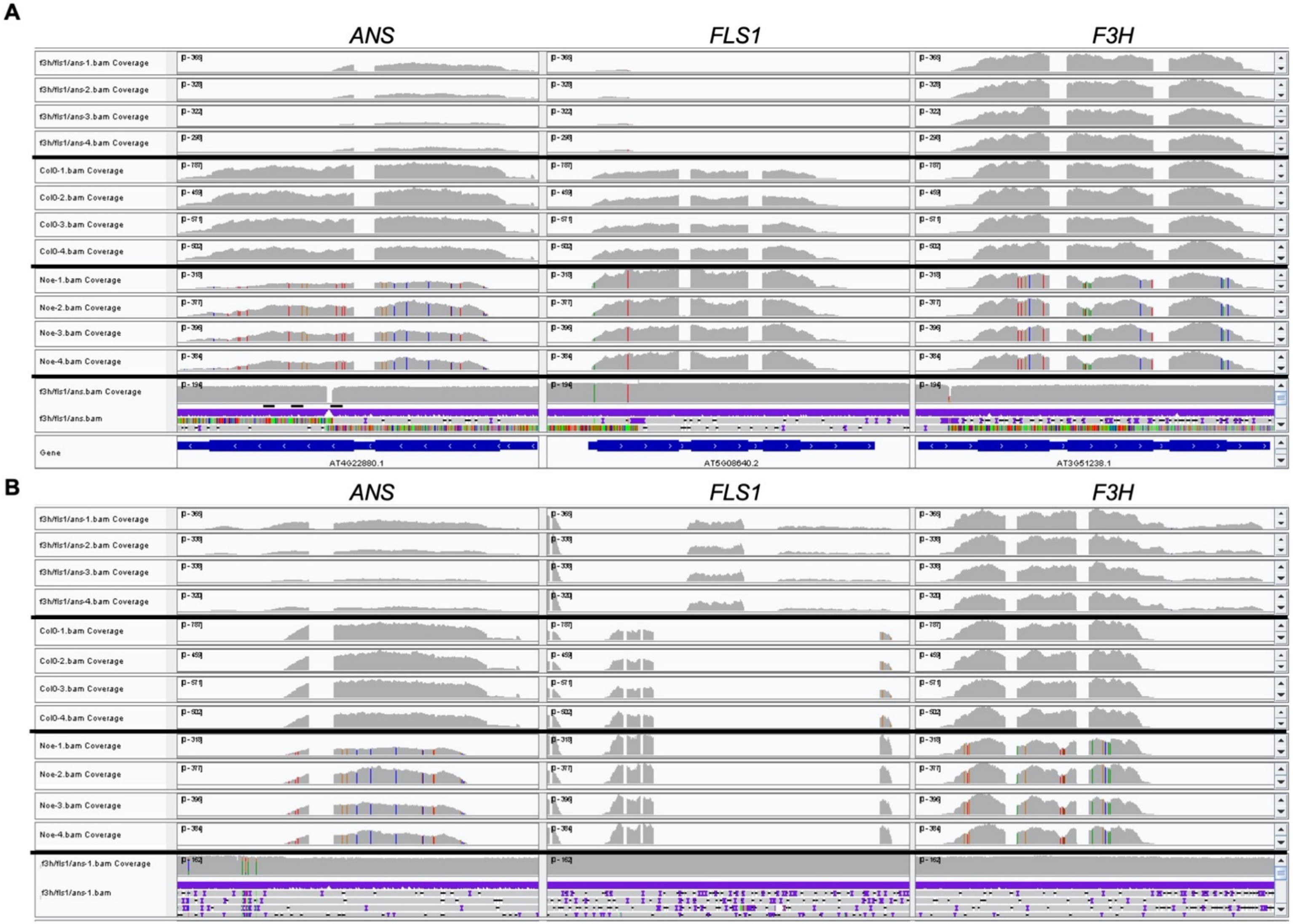


**Supplementary Figure S2: Genomic and transcriptomic characterisation of the mutant loci of the *A. thaliana f3h/fls1/ans* mutant.** (A) RNA-Seq read mappings against the TAIR10 Col-0 reference genome sequence. The four RNA-Seq mappings, corresponding to the four biological replicates of the *f3h/fls1/ans* mutant, are shown in row 1-4, while those of Col-0 follow in row 5-8 and Nössen in row 9-12. The 13th row shows the T-DNA and transposon insertion events in the *f3h/fls1/ans* mutant corresponding to the ONT read mapping of the mutant to TAIR10. The last row contains the gene structure based on the TAIR10 annotation. (B) Read mappings against the *f3h/fls1/ans* assembly. The same numeration as in panel A applies. However, in the last row the read mapping of the ONT reads against the *f3h/fls1/ans* assembly is shown.

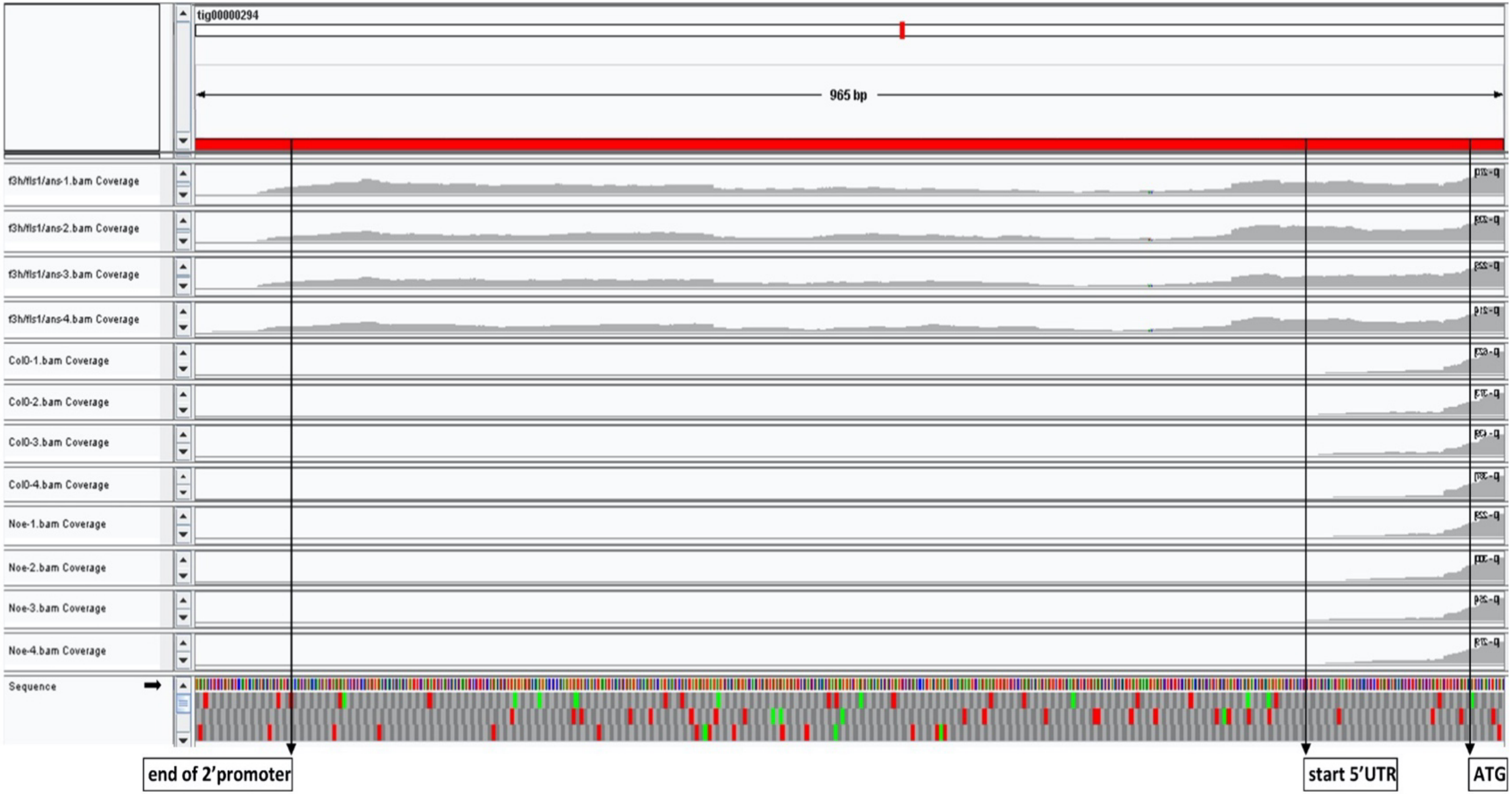


**Supplementary Figure S3: The *f3h* locus of the *A. thaliana f3h/fls1/ans* mutant.** RNA-Seq read mappings against the *f3h/fls1/ans* assembly. The four RNA-Seq read mappings corresponding to the four biological replicates of the *f3h/fls1/ans* mutant are shown in row 1-4, while those of Col-0 follow in row 5-8 and Nössen in row 9-12. The last row shows the three reading frames. Per frame the green boxes represent the position of potential start codons. Red boxes mark stop codons. Additionally, the end of the 2'promoter of the T-DNA, the start of the *F3H* 5'UTR and the typically used start codon are marked with black arrows and respective descriptions.

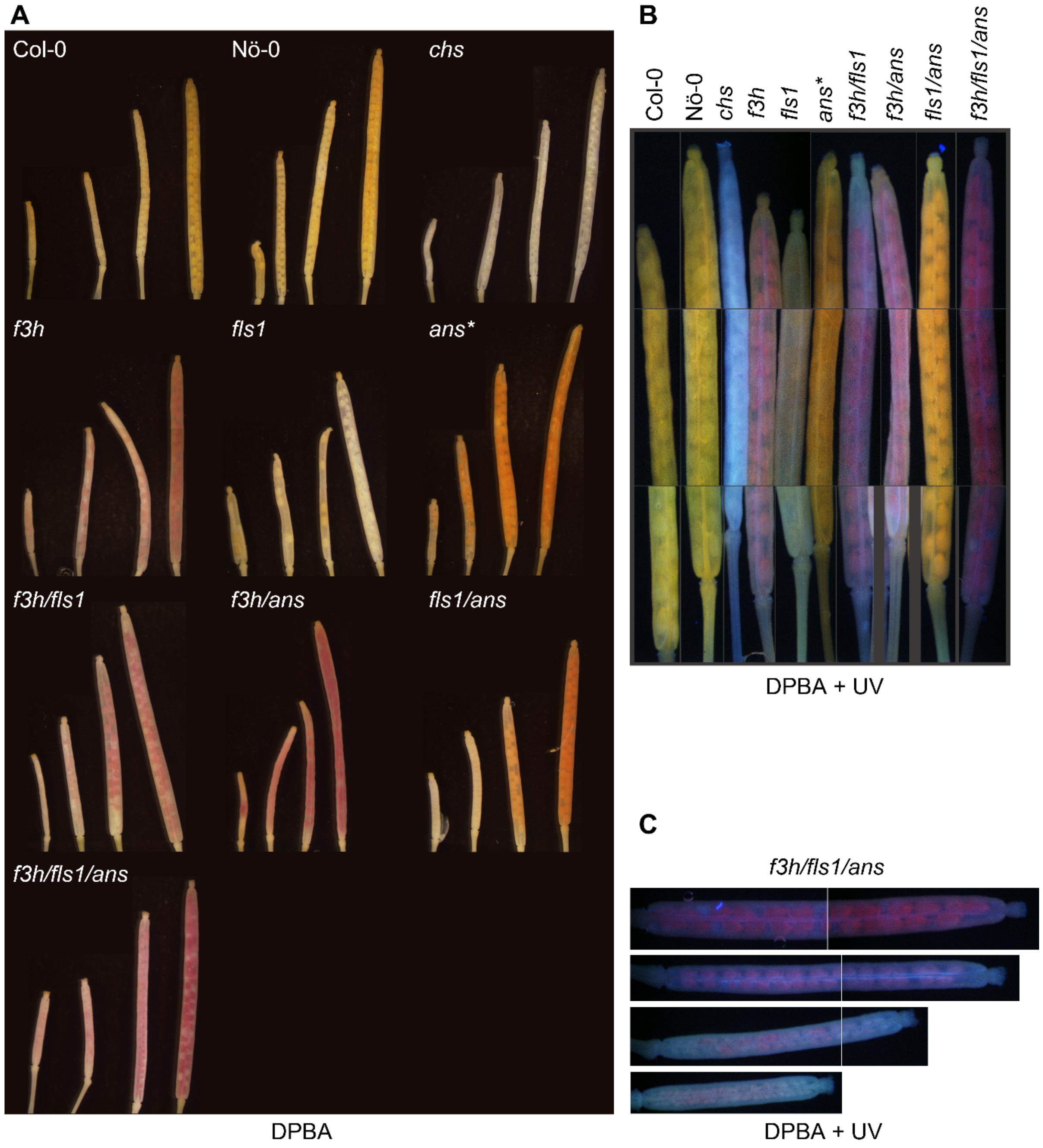


**Supplementary Figure S4: *In situ* flavonoid staining of siliques from *A. thaliana* wildtype and 2-ODD mutant plants.** Representative pictures of flavonoid accumulation in ethanol bleached and diphenylboric acid 2-aminoethyl ester (DPBA)-stained siliques under (A) white light and (B) UV light. (C) Close-up of different ages *f3h*/*fls1*/*ans* siliques under UV light. *please note: for the *ans* single mutant the *tt18-1* allele was used instead of *tds4-4.*

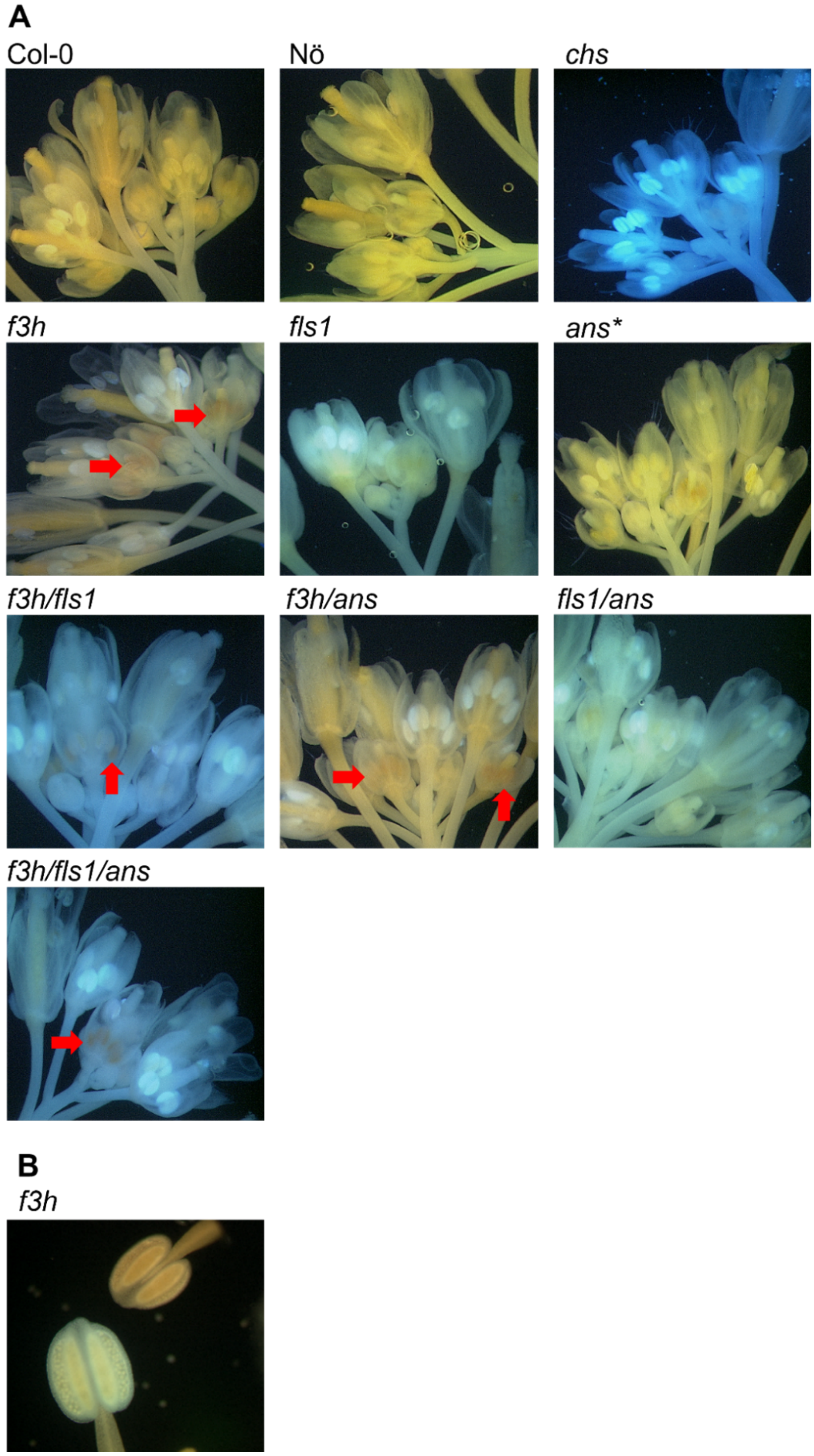


**Supplementary Figure S5: *In situ* flavonoid staining of inflorescences from *A. thaliana* wildtype and 2-ODD mutant plants.** Representative pictures of flavonoid accumulation in ethanol bleached and diphenylboric acid 2-aminoethyl ester (DPBA)-stained (A) inflorescences and (B) *f3h* stamen under UV light. Red arrows mark the presence of red fluorescence. *please note: for the *ans* single mutant the *tt18-1* allele was used instead of *tds4-4.*

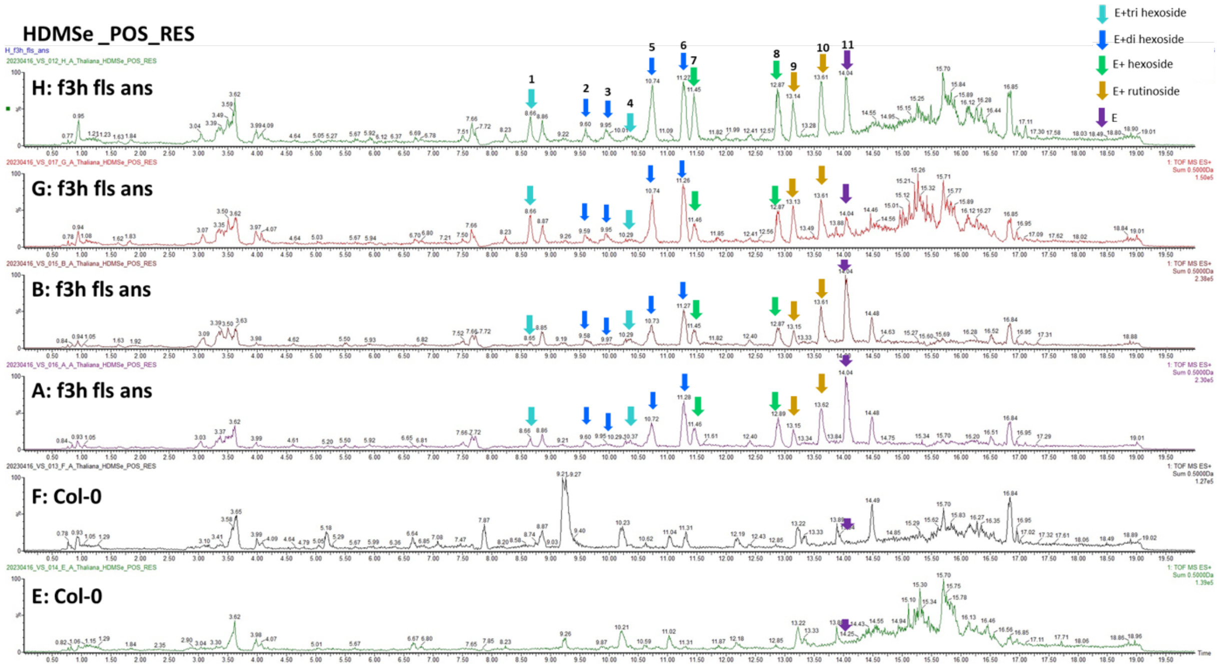


**Supplementary Figure S6: Comparison of the eriodictyol profile between the *A. thaliana f3h/fls1/ans* mutant and Col-0 in UPLC-HDMS$^{e}$ ESI+.** Peak numbers correspond to those given in Supplementary Table S4. E, eriodictyol.

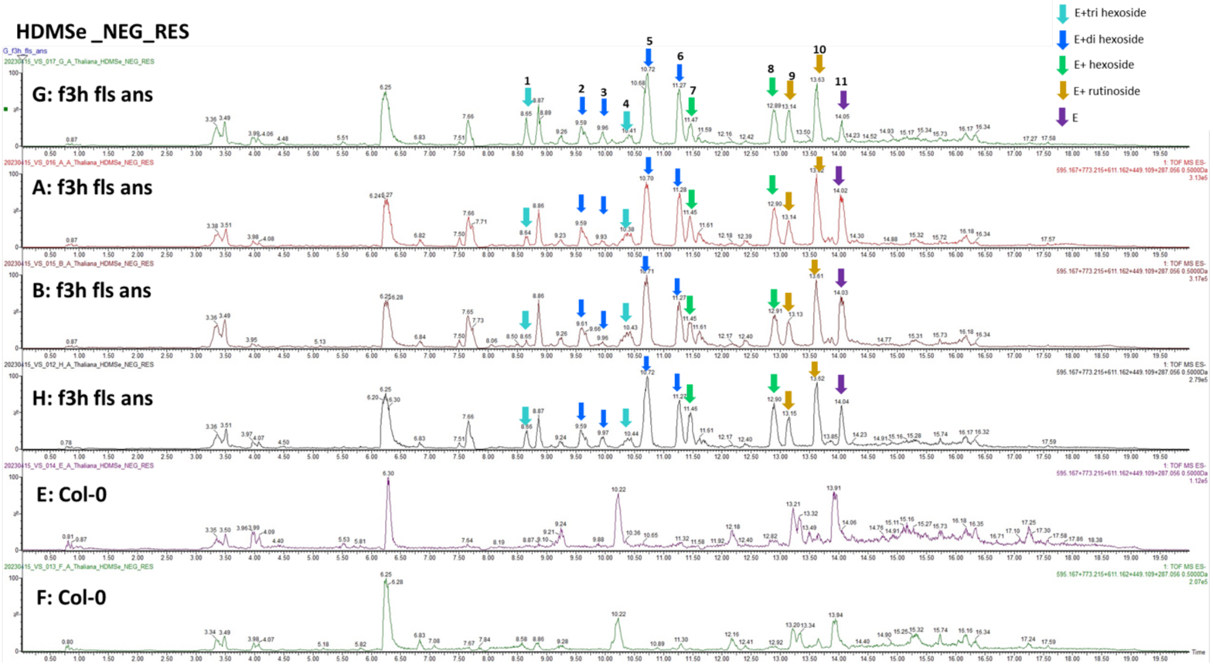


**Supplementary Figure S7: Comparison of the eriodictyol profile between the *A. thaliana f3h*/*fls1*/*ans* mutant and Col-0 in UPLC-HDMS$^{e}$ ESI-.** Peak numbers correspond to those given in Supplementary Table S4. E, eriodictyol.

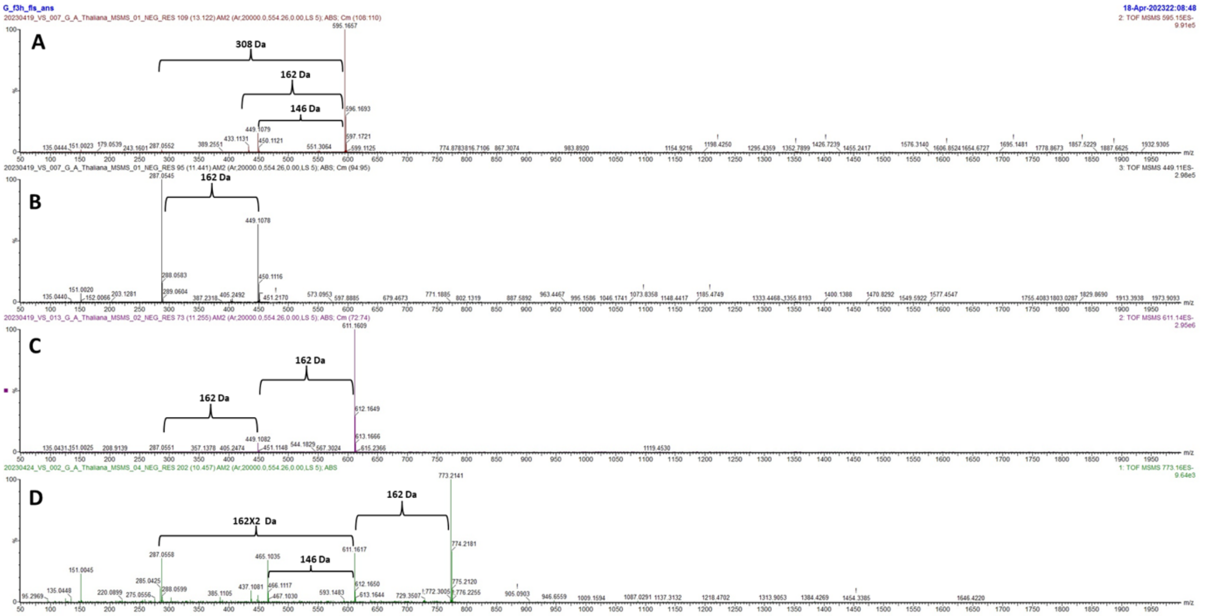


**Supplementary Figure S8: UPLC-MS/MS profile of one biological replicate from the *A. thaliana f3h*/*fls1*/*ans* mutant in ESI- mode.** Eriodictyol rutinoside or neohesperidoside (A), eriodictyol-7-*O*-glucoside (B), eriodictyol di-hexoside (C), eriodictyol tri-hexoside (D).

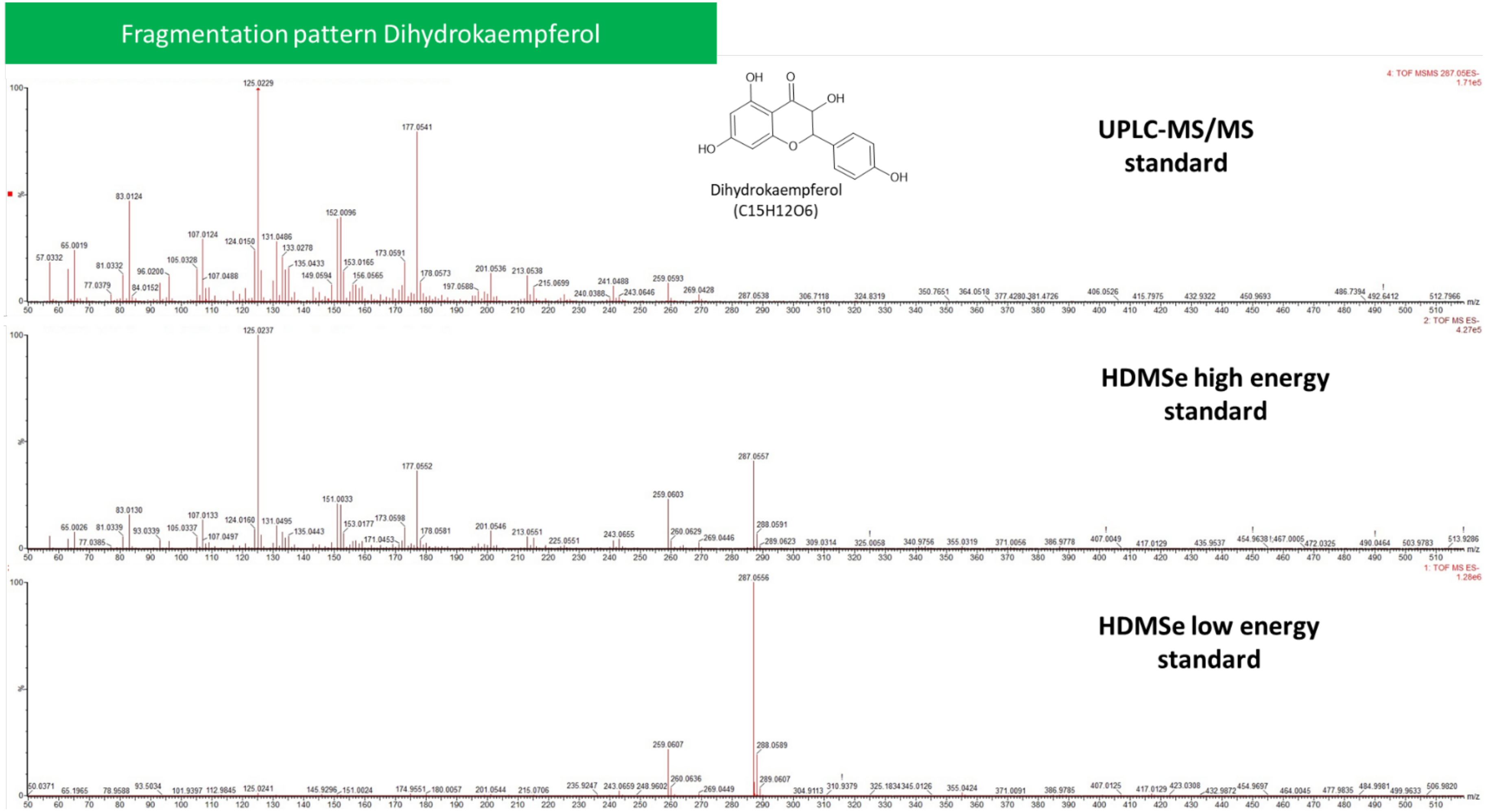


**Supplementary Figure S9**: ESI- fragmentation of dihydrokaempferol standard by UPLC-MS/MS and UPLC-TWIM-QToF (HDMS$^{e}$) high resolution mass spectrometry.

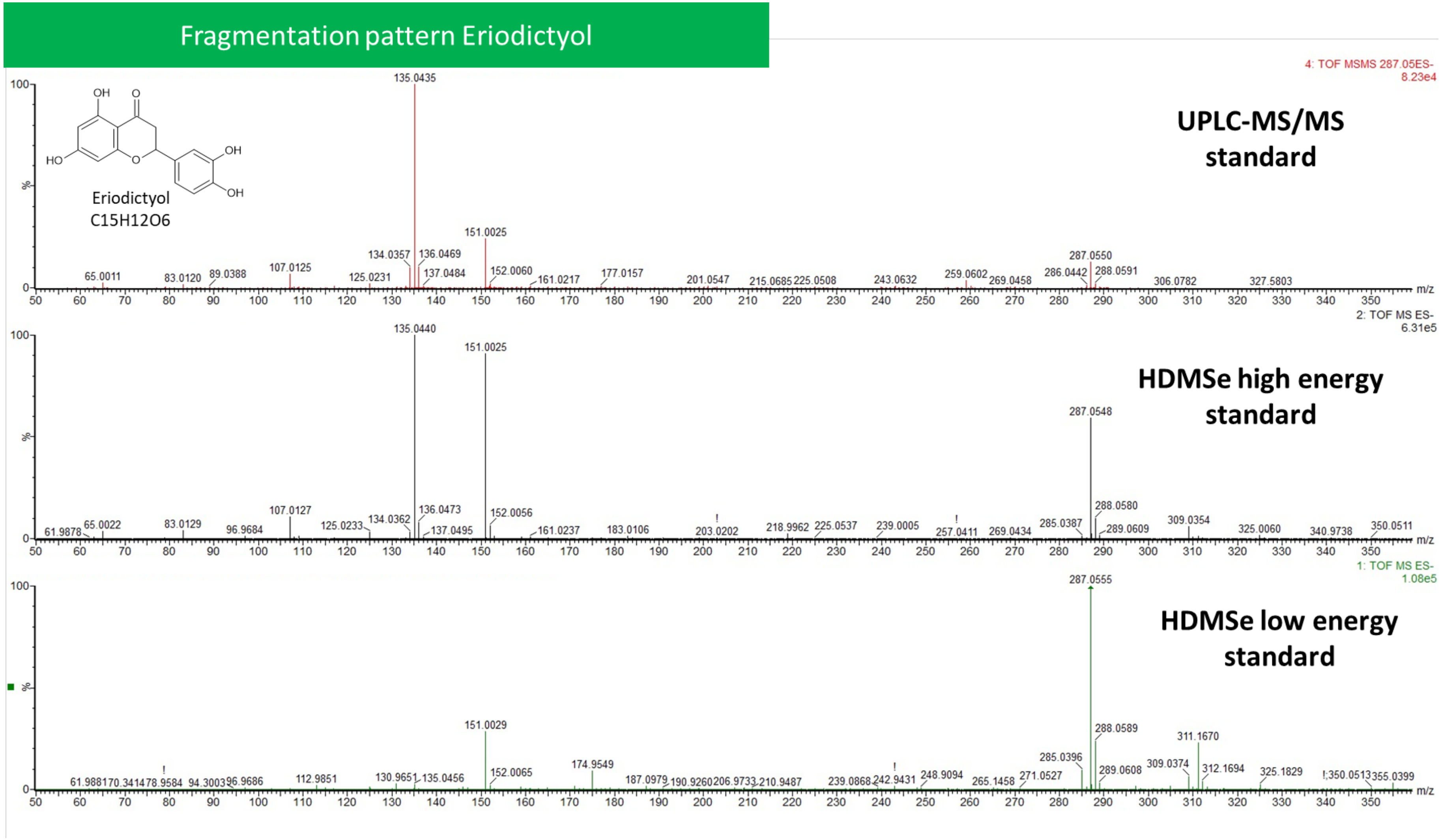


**Supplementary Figure S10**: Fragmentation of eriodictyol standard by UPLC-MS/MS and UPLC-TWIM-QToF (HDMS$^e$) high resolution mass spectrometry.

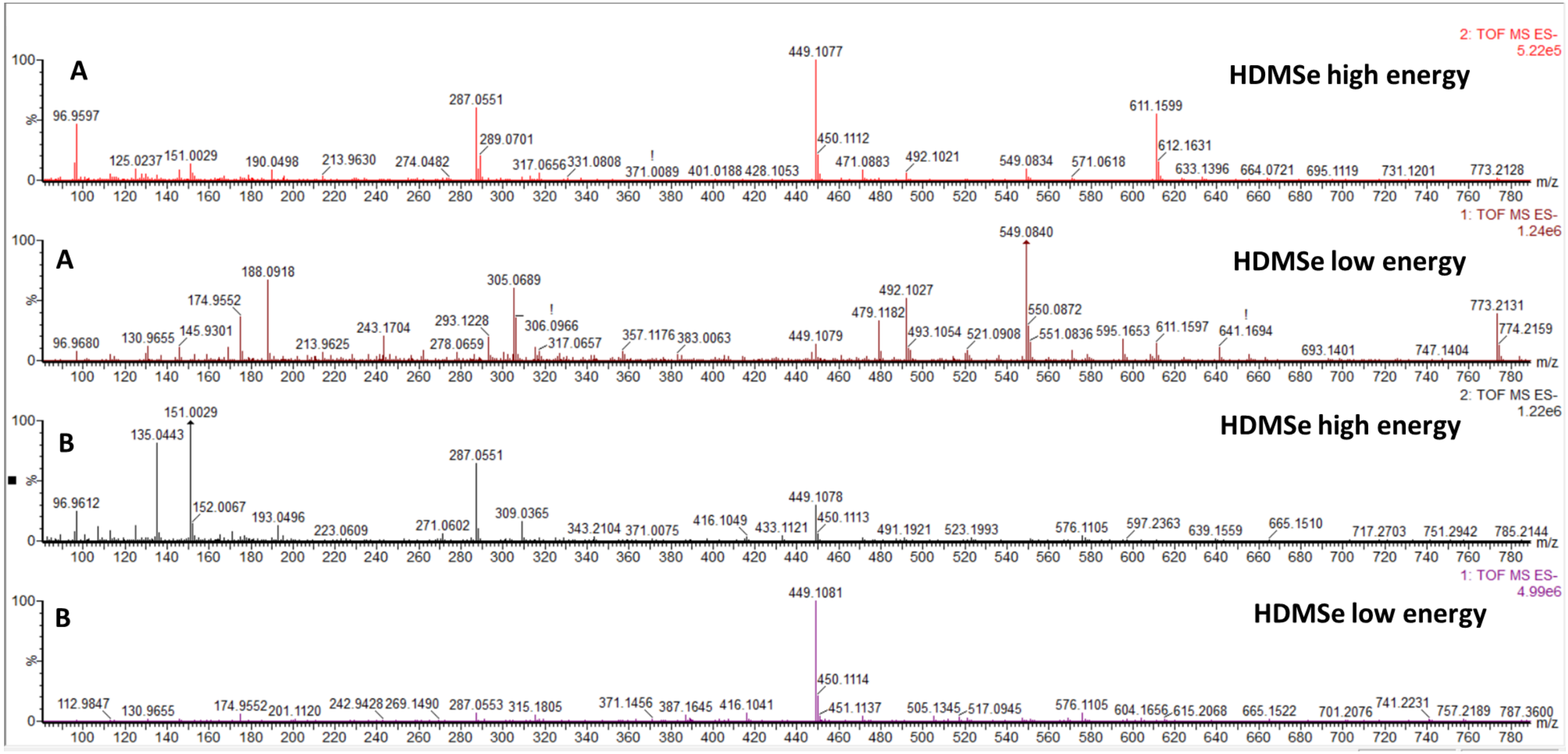


**Supplementary Figure S11:** ESI- UPLC-TWIM-QToF (HDMS$^{e}$) high resolution mass spectrometry of compound 1 and 8 (Supplementary Table S4) in the *f3h/fls1/ans* triple mutant biological replicate.

*F3'H*

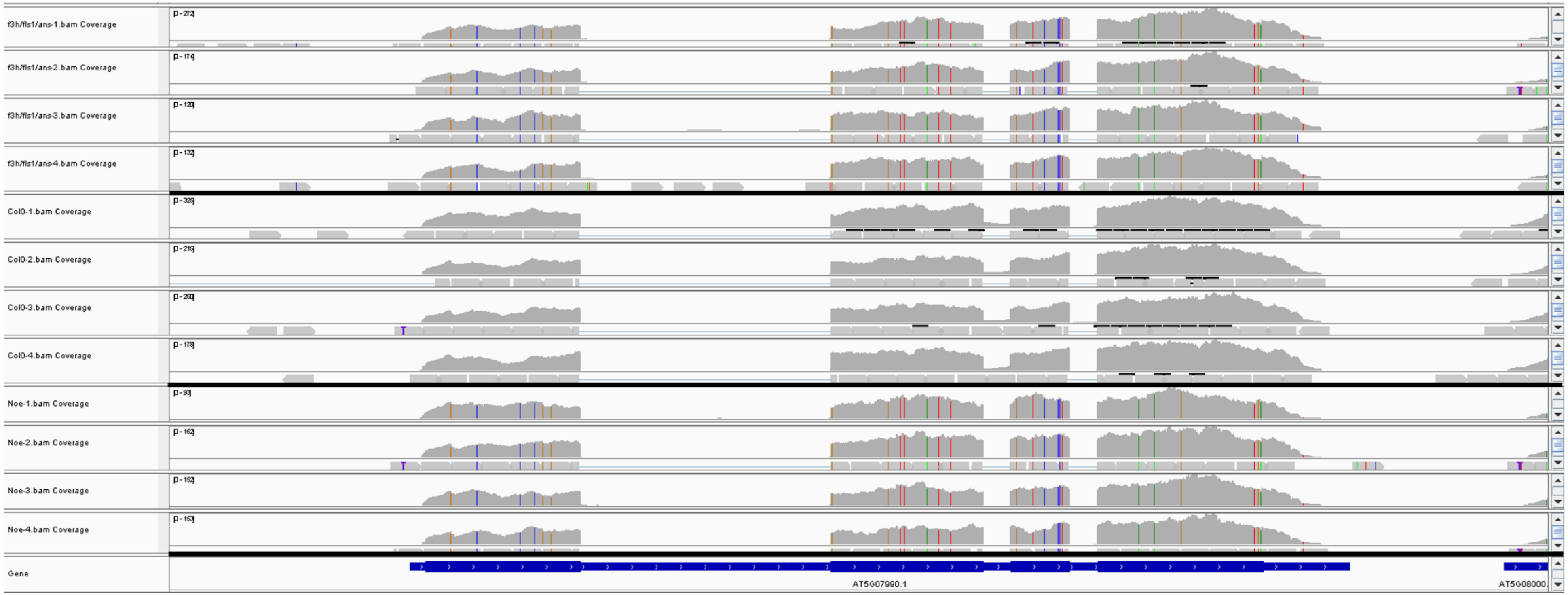


**Supplementary Figure S12: The *F3'H* locus of the *A. thaliana f3h/fls1/ans* mutant.** (A) RNA-Seq read mappings against the TAIR10 Col-0 reference genome sequence. The four RNA-Seq mappings, corresponding to the four biological replicates of the *f3h/fls1/ans* mutant, are shown in row 1-4, while those of Col-0 follow in row 5-8 and Nössen in row 9-12. The last row contains the gene structure based on the TAIR10 annotation.
