## Supplementary Table S1 for "Generation and characterisation of an *Arabidopsis thaliana f3h*/*fls1*/*ans* triple mutant that accumulates eriodictyol derivatives"

**Supplementary Table S1: *A. thaliana f3h/fls1/ans* mutant *de novo* ONT genome assembly statistics.**

| Parameter | Value |
|---|---|
| Number of contigs | 162 |
| Maximal contig length | 16,321,703 bp |
| Assembly size | 160,571,748 bp (160.6 Mbp) |
| GC content | 39.11 % |
| N50 | 8,170,251 bp |
| N90 | 246,021 bp |
| BUSCOs | 95.5% complete BUSCOs<br>[Single: 88.3%. Duplicated: 7.2%]; Fragmented: 1.7%. Missing: 2.8% |

Abbreviations: ONT, Oxford Nanopore Technologies; BUSCOs, Benchmarking Universal Single-Copy Orthologs

**Supplementary Table S2: Oligonucleotide primers used in this work.**

| Gene | Name | Sequence (5'-3') | Description |
|---|---|---|---|
| *F3H* | MM70 | GAGGTGACTAGAGACGATGAATCC | genotyping primer for the *F3H* wildtype and *tt6-2* mutant allele |
| | I250 | AATGACGATTGGGATTTTTGTAAC | genotyping primer for the *F3H* wildtype allele |
| | 8409 | ATATTGACCATCATACTCATTGC | GABI-KAT T-DNA LB primer |
| *FLS* | UH9 | CACTGAGATCTGTATGAGCCGGTACACC | genotyping primer for the *FLS* wildtype and *fls1-2* mutant allele |
| | RSt647 | TTACACATATCAACACGTACTTTA | genotyping primer for the *FLS* wildtype allele |
| | RSt645 | CCGGATCGTATCGGTTTTCG | RIKEN Ds-transposon primer |
| *ANS* | B078 | TTCCCCTGTTTTTAAGTTTATTT | genotyping primer for the *ANS* wildtype and *tds4-4* mutant allele |
| | B079 | AGAAAGACACAAACACATTATAAA | genotyping primer for the *ANS* wildtype allele |
| | RS630 | GCGTGGACCGCTTGCTGCAACTCTCTCAGG | SALK T-DNA LB primer |
